# A draft genome sequence of the miniature parasitoid wasp, *Megaphragma amalphitanum*

**DOI:** 10.1101/579052

**Authors:** Artem V. Nedoluzhko, Fedor S. Sharko, Brandon M. Lê, Svetlana V. Tsygankova, Eugenia S. Boulygina, Sergey M. Rastorguev, Alexey S. Sokolov, Fernando Rodriguez, Alexander M. Mazur, Alexey A. Polilov, Richard Benton, Michael B. Evgen’ev, Irina R. Arkhipova, Egor B. Prokhortchouk, Konstantin G. Skryabin

**Affiliations:** Nord University, Faculty of Biosciences and Aquaculture, Bodø, 8049, Norway.; National Research Center “Kurchatov Institute”, Moscow, 123182, Russia; Institute of Bioengineering, Research Center of Biotechnology of the Russian Academy of Sciences, Moscow, 117312, Russia; Josephine Bay Paul Center for Comparative Molecular Biology and Evolution, Marine Biological Laboratory, Woods Hole, Massachusetts 02543; Lomonosov Moscow State University, Faculty of Biology, Moscow, 119234, Russia; Center for Integrative Genomics, Faculty of Biology and Medicine, Génopode Building, University of Lausanne, CH-1015 Lausanne, Switzerland; Institute of Molecular Biology RAS, Moscow, 119991, Russia

## Abstract

Body size reduction, also known as miniaturization, is an important evolutionary process that affects a number of physiological and phenotypic traits and helps animals to conquer new ecological niches. However, this process is poorly understood at the molecular level. Here, we report genomic and transcriptomic features of arguably the smallest known insect – the parasitoid wasp, *Megaphragma amalphitanum* (Hymenoptera: Trichogrammatidae). In contrast to expectations, we find that the genome and transcriptome sizes of this parasitoid wasp are comparable to other members of the Chalcidoidea superfamily. Moreover, the gene content of *M. amalphitanum* compared to other chalcid wasps is remarkably conserved. Among the very rare cases of apparent gene loss is *centrosomin*, which encodes an important centrosome component; the absence of this protein might be related to the large number of anucleate neurons in *M. amalphitanum.* Intriguingly, we also observed significant changes in *M. amalphitanum* transposable element dynamics over time, whereby an initial burst was followed by suppression of activity, possibly due to a recent reinforcement of the genome defense machinery. Thus, while the *M. amalphitanum* genomic data reveal certain features that may be linked to the unusual biological properties of this organism, miniaturization is not associated with a large decrease in genome complexity.

## INTRODUCTION

Miniaturization in animals is an evolutionary process that is frequently accompanied by structural simplification and size reduction of organs, tissues and cells [1, 2]. The parasitoid wasp *Megaphragma amalphitanum* (Hymenoptera: Trichogrammatidae, subfamily Oligositinae) is one of the smallest known insects, whose size (250 μm adult length) is comparable with unicellular eukaryotes and even some bacteria (**Figure 1**). Parasitoids from the genus *Megaphragma* parasitize greenhouse thrips *Heliothrips haemorrhoidalis* (Thysanoptera: Thripidae) developing on the shrubs *Viburnum tinus* (Adoxaceae) and *Myrtus communis* (Myrtaceae) [3], and possibly *Hercinothrips femoralis* (Thysanoptera: Thripidae) [4]. The wasp spends most of its life cycle in host eggs, while the imago stage is very short and lasts only a few days [3, 4]. *M. amalphitanum* belongs to chalcid wasps, which represent one of the largest insect superfamilies (~23,000 described species)[5]. The higher-level taxonomic relationships of Trichogrammatidae, Chalcidoidea and Hymenoptera have been investigated in several recent studies [6–10] that helped to establish the placement of this unique taxon that related to Mymaridae and Pteromalidae.

**Figure 1.**
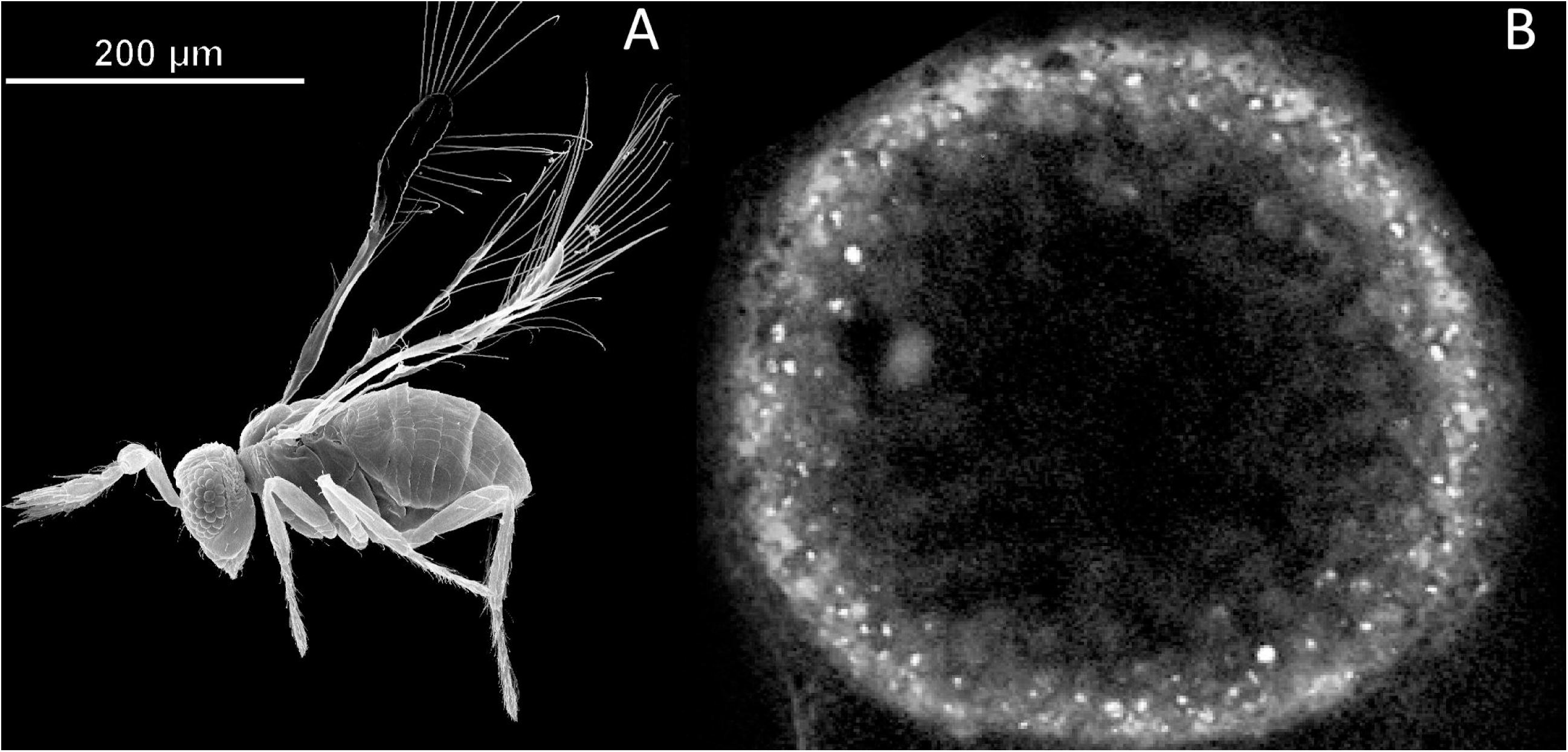
Size comparison of the parasitoid wasp *M. amalphitanum* and bacterium *Thiomargarita namibiensis*. (A) An adult stage of the parasitoid wasp *M. amalphitanum* (image adapted from [5]), (B) *T. namibiensis* – the largest known bacterium (modified from Schulz et al. 1999) [11].

Amongst notable anatomical features of *M. amalphitanum*, this species has only ~4600 neurons in its brain, which is substantially fewer than in the brains of other wasps, e.g. the parasitoid chalcid wasp *Trichogramma pretiosum* (Trichogrammatidae: Trichogrammatinae) (~18,000 neurons), *Hemiptarsenus sp.* (Chalcidoidea: Eulophidae) (~35,000 neurons), and the honey bee *Apis mellifera* (Apidae) (~850,000-1,200,000 neurons). Moreover, by the final stage of *M. amalphitanum* development, up to 95 percent of the neurons of the central nervous system lose their nuclei [12, 13]. Nevertheless, adult wasps, which have an average lifespan of 5 days, still preserve the basic functional traits of hymenopteran insects including flight, mating and oviposition in hosts [14].

In this study, we present a draft *M. amalphitanum* genome, and adult transcriptome, and compare these with several parasitoid wasp species of different body sizes from the Chalcidoidea and Ichneumonoidea hymenopteran superfamilies. We performed a comprehensive analysis of its coding potential, including both general gene ontology and pathway analyses as well as specific gene categories of interest, such as chemosensory receptors, venom components etc. Additionally, we investigated transposable element (TE) content and dynamics across several parasitoid wasp species and analyzed the major components of the genome defense machinery. As body size reduction and loss of physiological or phenotypic traits is often correlated with genome size diminution [15, 16] and/or gene networks reduction [17], including chromatin diminution from the somatic tissues during embryogenesis[18, 19], we initially anticipated that the *M. amalphitanum* genome would be greatly simplified during miniaturization.

## MATERIAL AND METHODS

### Detailed information is presented in ***Supplementary Information***

#### Nucleic acid extraction and library construction

*M. amalphitanum* individuals were reared in the laboratory conditions from eggs of *Heliothrips haemorrhoidalis* (Thysanoptera: Thripidae) collected in Santa Margherita, Northern Italy. DNA was extracted from ten individuals (males and females) using NucleoSpin Tissue XS kit (Macherey-Nagel, Germany) for each DNA-library. Three DNA libraries (DNA-library1 – whole insects; DNA-library2 – thorax and abdomen; DNA-library3 – head) were constructed using Ovation Ultralow Systems V2 kit (NuGEN, USA). Limited amount of biological material and low quantity of starting material (1 – 3 ng) did not permit construction of mate-paired libraries. Genome was sequenced used Illumina HiSeq 1500 (Illumina, USA) with 150 bp paired-end reads. RNA was extracted from ten *M. amalphitanum* individuals (males and females) using the Trizol reagent (Thermo Fisher Scientific, USA) by a standard protocol, and cDNA libraries were constructed using Ovation RNA-Seq System V2 kit (NuGEN, USA) with poly(A) enrichment. *Genome de novo assembly.* The output from Illumina sequencing of the genomic DNA library (source format *.fastq) was used for *de novo* genome assembly. To assemble the complete genome of *M. amalphitanum*, we used 102,188,833 paired-end reads. Genome assemblies have been constructed using different assembly algorithms, and their performance was compared to each other (**Figure S1**). Additionally, genomic DNA-libraries from thorax and abdomen (DNA-library2) of *M. amalphitanum* (SRR5982987) and from head (DNA-library3) of *M. amalphitanum* (SRR5982986) were prepared. Totally, 79,317,970 (paired-end sequencing: 2×100 bp) and 85,409,775 (single--end sequencing: 50 bp) DNA reads were sequenced and were used for *M. amalphitanum* coverage increase and as additional evidence during the search for missing genes (**Table S1**).

#### Transcriptome de novo assembly

Illumina RNA sequencing generated a total of 59,790,973 paired-end reads. Transcriptome *de novo* assembly was conducted using the default k-mer size in the Trinity software package (v. 2.4.0) [20], which combines three assembly algorithms: Inchworm, Chrysalis and Butterfly. Annotation of the *M. amalphitanum* transcriptome assembly was performed using the Trinotate pipeline (https://trinotate.github.io/).

#### Transposable element (TE) de novo identification and analysis

For *de novo* TE library construction, we used the REPET package [21] which combines three mutually complementing repeat identification tools (RECON, GROUPER and PILER), yielding a combined repeat library with the average consensus sequence length of 1.66 kb (ranging from 157 to 14,640 bp). The outputs were subject to additional classification with the RepeatClassifier tool from the RepeatMasker package (www.repeatmasker.org), which was also used to build the corresponding TE landscape divergence plots.

## RESULTS AND DISCUSSION

### Whole genome and transcriptome sequencing and assembly of *M. amalphitanum*

To gain insight into the genomic signatures of miniaturization that would distinguish *M. amalphitanum* from other Hymenoptera, we performed whole-genome shotgun sequencing of DNA (DNA-library1) isolated from ten adult individuals (males and females), using the Illumina platform (**Table S1**). The resulting genome assembly (PRJNA344956) has a cumulative length of 346 megabases (Mb), with a scaffold N50 □ of 10,296 bp. The total genome coverage is 88.6-fold. Thus, the genome of *M. amalphitanum* is comparable in size with other Chalcidoidea wasps, such as *Copidosoma floridanum, T. pretiosum* or *Nasonia vitripennis* [22, 23]. The best-performing combination of assembly software yielded contig N50 of 4,285 bp and allowed us to assemble 94,687 scaffolds from the quite low amounts of starting DNA material (**Table S2-S3; Figure S1**).

The *M. amalphitanum* genome assemblies were evaluated with the BUSCO (benchmarking universal single-copy orthologs) arthropod gene set [24], which uses 2,675 near-universal single-copy orthologs to assess the relative completeness of genome assemblies. Through this analysis, 7.55% of the conserved genes were initially identified in the *M. amalphitanum* assembly as missing (**Table S4**). More detailed information on our extensive search for the missing genes in *M. amalphitanum* genome is presented below.

We also performed whole-body transcriptome analysis using RNA extracted from ten *M. amalphitanum* individuals (males and females). Transcriptome *de novo* assembly (PRJNA344956) was performed using the Trinity software [20]. A total of 46,841 contigs were assembled with a mean length of 586 bp and an N50 of 633 bp from the quite low amounts of starting RNA material (**Table S5**). The Illumina paired-end RNA-Seq data from *M. amalphitanum* were mapped to the previously assembled genome using bowtie2 [25]. Inspection of the alignments revealed that 79.95% of reads could be mapped to the genome. The BUSCO statistics for the transcriptome assembly is also presented in **Table S4**.

### Gene ontology analysis

We used Gene Ontology (GO) analysis terms to describe characteristics of *M. amalphitanum* gene products in three independent categories: biological processes (**Figure S2**), molecular function (**Figure S3**), and cellular components (**Figure S4**). BLASTX outputs were used to retrieve the associated gene names and GO terms in all three categories (**Table 1**).

**Table 1.** Basic Gene Ontology (GO) analysis terms for *M. amalphitanum* gene products

| GO assignments of the transcripts | Transcript counts and percentage from total |  |  |
| --- | --- | --- | --- |
| Biological processes | 8,812 counts, 49.72% |  |  |
|  | Transcription | Regulation of transcription | DNA integration |
|  | 15 % | 10 % | 8 % |
| Cellular components | 4,802 counts, 27.10% |  |  |
|  | Nucleus and cytoplasm components | Integral membrane components | Plasma membrane components |
|  | 18 % | 9 % | 7% |
| Molecular functions | 4,108 counts, 23.18% |  |  |
|  | ATP binding | Metal ion binding | Zinc ion binding |
|  | 17 % | 12 % | 10 % |

All *M. amalphitanum* transcripts were matched to the Clusters of Orthologous Groups (COG) database to predict and classify their functions. In total, 8,810 genes were assigned to 25 COG functional categories. One of the largest groups is represented by the cluster for post-translational modification, protein turnover, and chaperones (988 counts; 10.7%), followed by intracellular trafficking, secretion, and vesicular transport (659 counts; 7.2%), DNA replication, recombination and repair (606 counts; 6.6%), signal transduction mechanisms (599 counts, 6.5%) and transcription (587; 6.4%) (**Figure S5**).

To better understand incorporation of genes into diverse pathways, all annotated transcripts were mapped against the KEGG database for pathway-based analysis. As a result, 6,130 transcripts out of a total of 46,841 were assigned to a KEGG pathway, and were present in 328 different KEGG pathways. The KEGG pathway distribution is summarized in **Figure S6**. The top five pathways are metabolism (479 counts; 7.8%), biosynthesis of secondary metabolites (150 counts; 2.4%), RNA transport (100 counts; 1.6%), biosynthesis of antibiotics (95 counts; 1.5%), spliceosome (94 counts; 1.5%).

The annotation of *M. amalphitanum* and the available transcriptome assemblies of other parasitoid wasps from the families Trichogrammatidae *(T. pretiosum*, a lepidopteran egg parasitoid) and Braconidae including *Cotesia vestalis* (a diamondback moth parasitoid), *Diachasma alloeum* (an apple maggot parasitoid) and *Fopius arisanus* (tephritid fruit fly parasitoid) were used for comparative analysis of the most represented gene functions in parasitoids. We also used transcriptome assemblies from the Agaonidae fig wasp, *Ceratosolen solmsi.* We found significant similarities between *M. amalphitanum, T. pretiosum* and *C. vestalis* major GO enrichment categories (**Figure S7-S9**). At the same time, a significant number of transcripts related to DNA integration relative to other parasitoid wasps was found in *D. alloeum* and *M. amalphitanum* (**Figure S7**) (see below). Complete information about reference datasets used for *M. amalphitanum* genome and transcriptome data analysis is shown in **Table S6**. The Trinotate statistics for annotation of *M. amalphitanum, C. solmsi, D. alloeum, F. arisanus, C. vestalis* and *T. pretiosum* transcriptome assemblies is presented in **Table S7**.

### Missing genes and missing or rapidly evolving gene clusters in the *M. amalphitanum* genome

We clustered gene orthologs and identified gene clusters for each hymenopteran taxa (Chalcidoidea: *M. amalphitanum, T. pretiosum, C. solmsi*, C. *floridanum*, and *N. vitripennis;* Ichneumonoidea: *D. alloeum* and *F. arisanus;* Apoidea: *A. mellifera)* using OrthoMCL [26]. The core gene set of all the hymenopteran species was composed of 6278 gene clusters, 122 gene clusters were unique to the chalcid clade. 262 gene clusters were not detected in any of the chalcids analysed (**Supplementary Dataset 3**; NCBI BioProject: PRJNA344956), but found in all the other hymenoptera, consistent with a similar recent analysis [27]. Our findings suggest that that these gene losses occurred in the last common ancestor of chalcids or point to the possibility of parallel genome evolution across these species. Interestingly, the missing/rapidly evolving genes include homologs of genes that have important roles in embryonic patterning and development in other insects (e.g., *krueppel-1, knirps* or *short gastrulation* [27]).

To determine whether miniaturization in *M. amalphitanum* is associated with gene loss, genomic data of six larger hymenopteran species *(T. pretiosum, C. vestalis, C. floridanum, F. arisanus, N. vitripennis*, and *N. giraulti)* – as well as the well-annotated genome of the honeybee *(A. mellifera)* as reference – were used (body sizes are presented in **Table S6**). We mapped the *M. amalphitanum* (DNA-library1), *T. pretiosum, C. vestalis, C. floridanum, F. arisanus, N. vitripennis, N. giraulti* DNA reads on the *A. mellifera* genome sequence (PRJNA13343, PRJNA10625) (**Figure S10-S11**), and detected 115 genes that were not represented by *M. amalphitanum* sequencing reads but were present in other parasitoid wasps. We next increased the coverage of the *M. amalphitanum* genome to 146.8-fold by adding the reads from additional libraries (DNA-library2 and DNA-library3) (**Table S1**) and observed the apparent absence of 114 of the 115 genes. An additional TBLASTX search identified 36 of these genes as present, yielding a total of 78 missing genes (**Table S8**). However, querying the *M. amalphitanum* genome with the corresponding amino acid sequences from the closest wasp ortholog (*N. vitripennis* or *T. pretiosum)* in TBLASTN searches reduced the number of missing genes to just five: centrosomin, phosphoglycerate mutase 5, phosphoglycerate mutase 5-2, 26S proteasome complex subunit DSS1, and mucin-1/nucleoporin NSP1-like. We detected short *M. amalphitanum* genome sequences encoding protein fragments (~8-23 amino acid residues) with some similarity to four of them, suggesting that they may be in the process of degeneration in this species. Despite a careful search, we were unable to find any homologous sequence related to *centrosomin (cnn)* gene either in the assembled genome or our cDNA libraries (**Figure S15**). Although *cnn* is regarded as rapidly evolving [28], sequence homology can be readily discerned and orthologs are present in every other insect, including the parasitoid *T. pretiosum*, suggesting that this is a *bona fide* absent gene. Globally, however, these analyses indicate that there has been little gene loss in *M. amalphitanum*

### Chemosensory genes in the *M. amalphitanum* genome

Chemosensory receptors are encoded by some of the largest gene families in insect genomes, reflecting their important and wide-ranging roles in detection of environmental odors and tastants. We asked how these gene families have evolved in *M. amalphitanum*, whose central and peripheral nervous systems are highly reduced [2, 14]. The highly divergent sequences of chemosensory receptors and relatively short genomic contig lengths available for *M. amalphitanum* precluded accurate annotation of full-length sequences in this species for the majority of loci. Nevertheless, comparison with chemosensory receptor repertoires of other insects allowed us to define probable orthologous relationships with receptors of known function in other species and obtain initial estimates of the size of each family.

The most deeply conserved family of chemosensory receptors in insects are the Ionotropic Receptors (IRs), which are distantly related to ionotropic glutamate receptors [29, 30]. IRs function in heteromeric protein complexes comprising more broadly-expressed co-receptors with selectively expressed “tuning” IRs that determines sensory specificity. We identified orthologs of each of the co-receptors (Ir8a, *Ir25a* (two paralogs), *Ir93a* and *Ir76b*), as well as four genes encoding tuning IRs related to acid-sensing receptors in other species. We also identified orthologs of IR68a, which functions in hygrosensation [31] and IR21a, which functions in cool temperature-sensing [32, 33]. Overall, the repertoire of IRs in *M. amalphitanum* is therefore very similar in size and content to that of *N. vitripennis [30].*

Insects possess a second superfamily of chemosensory ion channels – distinguished by a heptahelical protein structure – comprising Odorant Receptor (OR) and Gustatory Receptor (GR) subfamilies, which generally function in detection of volatile and non-volatile stimuli, respectively [34–37]. Similar to IRs, ORs function in heteromeric complexes of a conserved co-receptor (ORCO) and a tuning OR. We identified an *M. amalphitanum* ortholog of *Orco* and 83 additional Or-related sequences. We caution that many of these *Or* sequences are small fragments (often located near the end of the assembled contigs), so it is currently difficult to determine whether these are intact genes or pseudogenes. Within the GR repertoire, we identified genes encoding proteins related to GR43a, a sensor of both external and internal fructose [38], two others similar to other insect sugar-sensing GRs [39], and 25 additional *Gr* gene fragments. The sizes of these repertoires are smaller than in *N. vitripennis* (300 *Ors* (including 76 pseudogenes) and 58 *Grs* (including 11 pseudogenes) [40]), but similar to non-miniaturized parasitoid wasps *Meteorus pulchricornis* and *Macrocentrus cingulum* [41, 42]. However, precise comparison with the latter two species is difficult, as receptors in these wasps were identified from antennal transcriptomes, thereby representing only one of these insects’ chemosensory organs.

In sum, these analyses reveal that despite drastic nervous system reduction, *M. amalphitanum* has retained the conserved chemosensory receptors of larger wasps (and other insects), and appears to have numerous additional order- or species-specific receptors to allow detection of environmental chemical cues.

### Venom components in the *M. amalphitanum* transcriptomic data

Parasitoid wasps often use venom to modify the metabolism of their hosts; toxins and their known or presumed biological functions are described in various species [43]. We investigated the presence of homologs of *N. vitripennis* toxin constituents in *M. amalphitanum* and other parasitoid wasps *(Megastigmus spermotrophus, N. vitripennis, C. solmsi, T. pretiosum)*, using previously published venom data [44, 45] and the transcriptomes of chalcid wasps (**Table S6**). We identified 28 transcripts encoding putative venom proteins (**Figure 2; Table S9**); homologs of these are found in all investigated Chalcidoidea species (**Table 2**). Assuming that most of these candidates are truly conserved venom proteins among Chalcidoids, *M. amalphitanum* venom’s diversity does not seem to have been significantly affected by size reduction.

**Figure 2.**
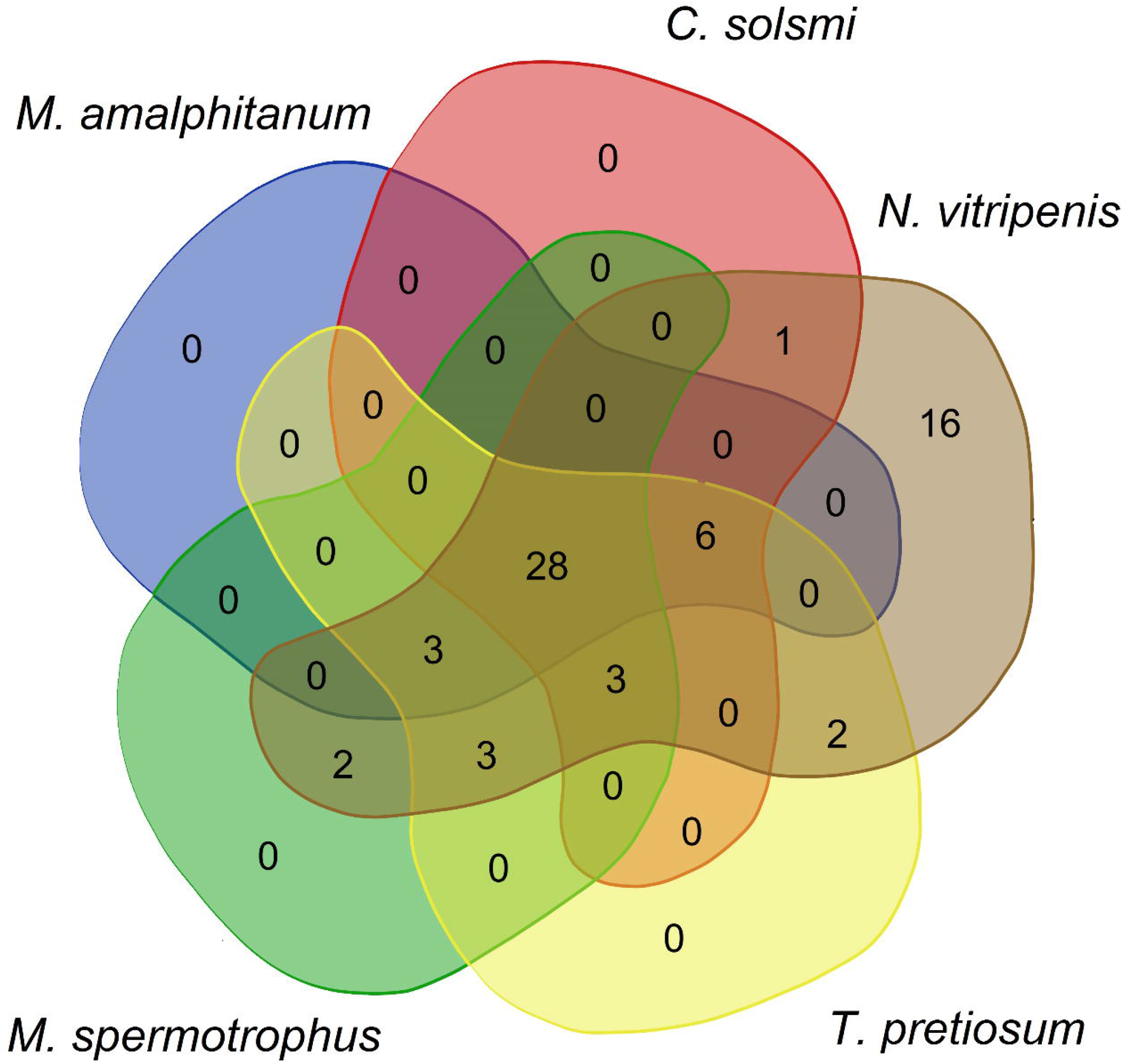
A Venn diagram showing *Nasonia vitripennis* venom components in other Chalcidoidea species: *M. spermotrophus, C. solmsi, T. pretiosum*, and *M. amalphitanum.*

**Table 2.** Number of homologs of *N. vitripennis* venom (*N. vitripennis* toxin constituents) in *M. amalphitanum* and other Chalcidoidea species based on Universal Chalcidoidea Database (http://www.nhm.ac.uk/our-science/data/chalcidoids/database/)

| Parasitoid wasp species | Families of Chalcidoidea | Number of *N. vitripennis* venom constituents | Body size, mm | Approximate number of hosts |
| --- | --- | --- | --- | --- |
| *M. amphitanum* | Trichogrammatidae | 37 | 0.25 | 2 insect species from one order |
| *C. solmsi* | Agaonidae | 38 | 2.7 | 2 plant species from one family |
| *M. spermotrophus* | Torymidae | 41 | 2.8 | 13 plant species from one family |
| *T. pretiosum* | Trichogrammatidae | 45 | 0.5 | >140 insect species from 4 orders |
| *N. vitripennis* | Pteromalidae | 64 | 2.2 | >110 insect species from 8 orders |

### *M. amalphitanum* transposable elements and genome defense

Transposable elements (TEs) constitute a measurable fraction of virtually all eukaryotic genomes, and can play important roles in their function and evolution. In insects, TE activity has been implicated in evolution of eusociality, based on comparison of ten bee genomes with increasing degrees of social complexity [46]. We performed *de novo* TE identification and comparative analysis of TE dynamics in *M. amalphitanum* and in a representative set of larger wasp genomes for which TE content has previously been reported: the parasitoid *N. vitripennis* and two primitively eusocial aculeate wasps *Polistes canadensis* and *Polistes dominula* [12, 23, 47]. Additionally, we analyzed TEs in the genomes of parasitoid wasps *T. pretiosum* from the family Trichogrammatidae and *D. alloeum* from the family Braconidae, in which TE content has not been studied.

For uniformity of measurements, we applied the same workflow to all genomes, without relying on pre-existing repeat libraries. We employed the REPET package for *de novo* TE identification (also used in [46]), and RepeatMasker for repeat classification and construction of TE landscape divergence plots. Comparison of the overall repeat content across six wasp species did not reveal substantial differences between four species (18.5% in *M. amalphitanum vs.* 18.1%, 17.7% and 14.2% in *P. canadensis, P. dominula* and *T. pretiosum*, respectively), while the *N. vitripennis* genome was found to be 32.5% repetitive, in close agreement with the published estimate [23], and *D. alloeum* was highly repetitive at 52.8% (pie charts in **Figure 3; Figure S12**). Surprisingly, TE dynamics over time, which is shown on the corresponding TE landscape divergence plots, was found to differ substantially for *M. amalphitanum*, which displayed a pronounced decline in recent TE activity after an initial increase, a pattern that was not observed in any other wasp (**Figure 3**).

**Figure 3.**
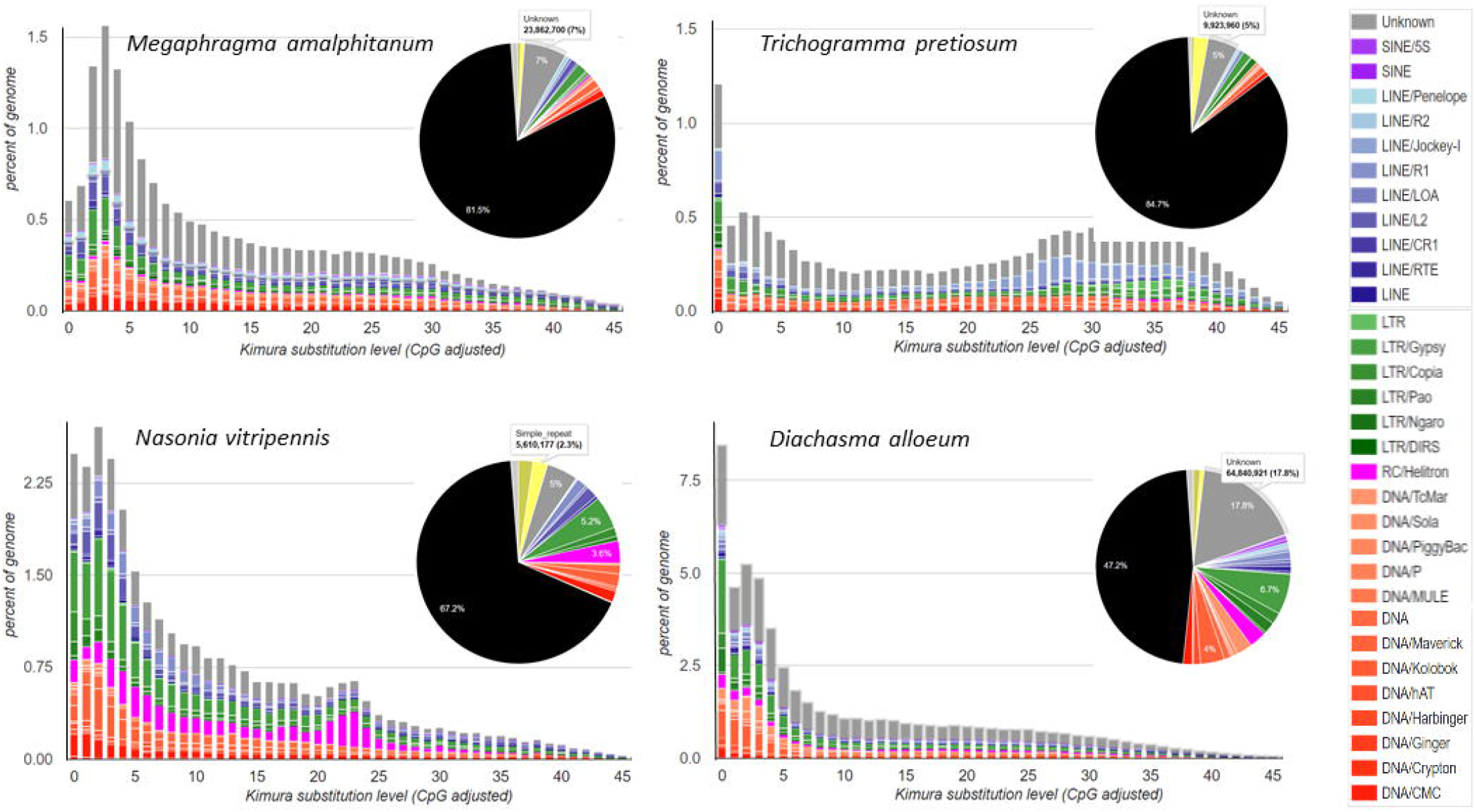
Comparison of TE landscape divergence plots and TE genome fraction pie charts in four parasitoid wasp species: *M. amalphitanum, T. pretiosum, N. vitripennis*, and *D. alloeum*.

While TE dynamics may be affected by different factors, the observed drop in active TE content in *M. amalphitanum* may be relevant to the unique biology of this highly miniaturized insect. Its closest relative, *T. pretiosum*, is about 2-fold larger in body length. The spike in recent TE activity in *T. pretiosum* may have been caused by *Wolbachia* infection, which typically results in abandonment of sexual reproduction [48] and the concomitant TE proliferation in a non-recombining genome [49]. Other wasps do not display notable drops or spikes in current TE activity; TE inactivation was reported in two asexual mites [50], however it appears to be ancient and may have occurred prior to the abandonment of sex. Overall, the continued decline in *M. amalphitanum* TE activity over the span of several million years – not observed in *T. pretiosum* which shares the most recent common ancestor with *M. amalphitanum* – represents a highly unusual genomic feature in every hymenopteran we examined, including ants (not shown). Interestingly, no traces of *Wolbachia* infection or other representatives of the Rickettsiaceae family were found in *M. amalphitanum* individuals [51], while the sequenced *T. pretiosum* carries the *Wolbachia* symbiont [52]; the sequenced *Nasonia* strain was maintained on antibiotics to cure it of infection.

To gain insights into possible reasons for reduction in TE activity after the initial burst, we investigated the major components of the genome defense machinery in *M. amalphitanum*, including Dicer (Dcr)-like and Argonaute (Ago)/Piwi-like protein-coding genes. In insects, *Ago-1* and *Dcr-1* homologs represent the key components of the miRNA pathway; *Ago-2* and *Dcr-2* mediate antiviral RNA interference; and *Piwi* and *Ago-3/Aub* suppress TE activity in the germline [53]. Both *M. amalphitanum* and *T. pretiosum* possess equal numbers of *Dcr-1* and *Dcr-2* homologs, as well as *Ago-2* and *Ago-3* homologs (**Figure S13**). However, in *M. amalphitanum*, the *Ago-1* and the *Piwi/Aub* homologs underwent a relatively recent duplication in comparison to *T. pretiosum* (**Figure 4**). This may indicate additional layers of enforcement in the miRNA and piRNA pathways of *M. amalphitanum*, both of which should result in suppression of TE activity.

**Figure 4.**
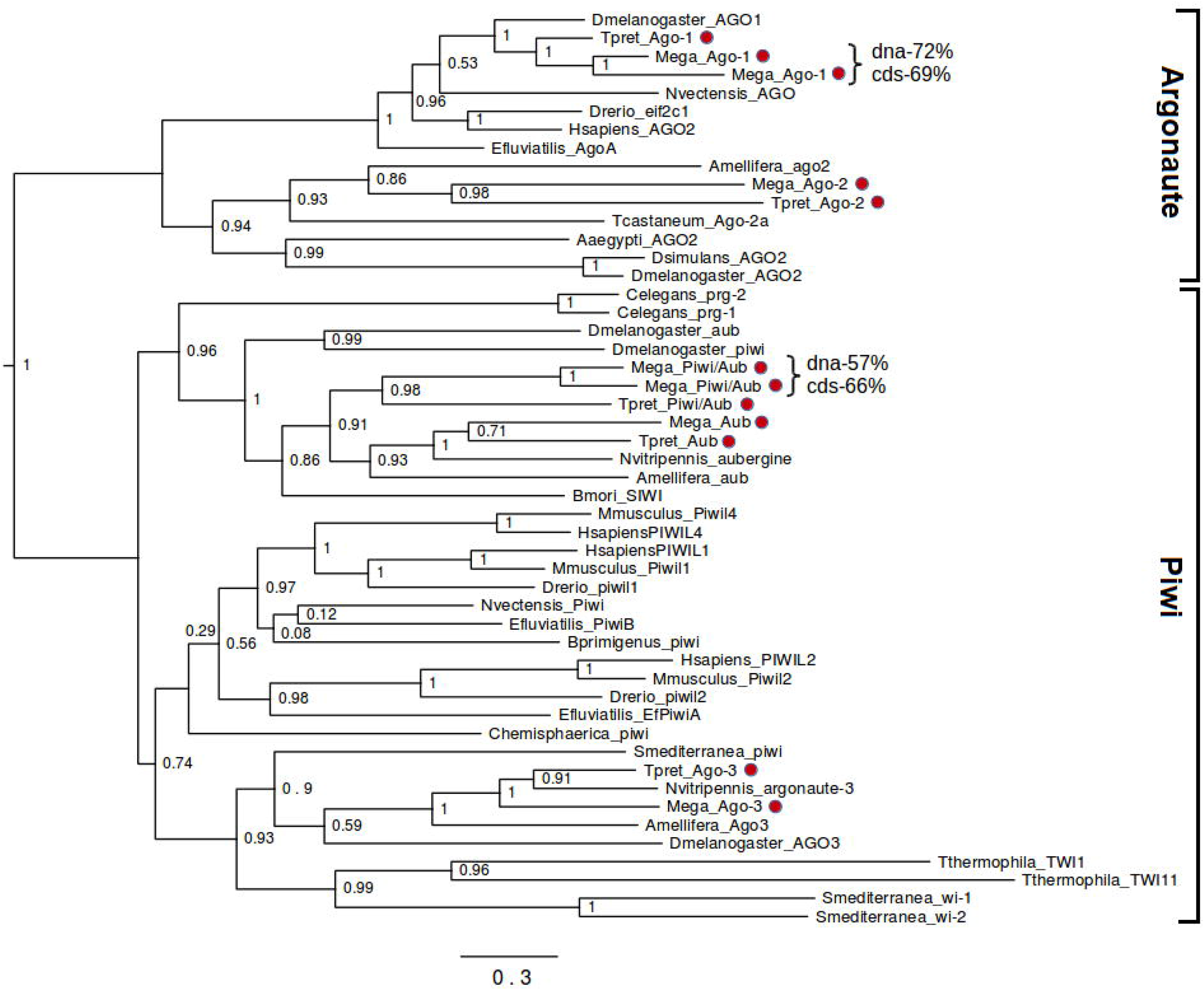
Maximum likelihood analysis of phylogenetic relationships between Piwi/Argonaute coding sequences. Percent identity is indicated for the duplicated Ago-1 and Piwi/Aub homologs, both at the DNA and protein (cds) level. Phylogeny analysis and notations are as in Supplementary Figure S13.

The drop in TE activity is also evident from the transcriptome analysis. The GO radar plot (**Figure S7**) shows a substantial number of short contigs related to DNA integration, most of which upon inspection were found to represent separate fragments of gypsy-like and copia-like LTR retrotransposons, and a few belong to Polinton, P and Ginger DNA TEs. Transcriptionally active copies fall into two groups: first, those which apparently proliferated during the burst of TE activity and have since accumulated debilitating mutations making them incapable of transposition, but still retain a certain level of transcriptional activity; second, those that originate from recent infections by retrovirus-like TEs and contain uninterrupted ORFs, but are not actively proliferating and are present at very few genomic loci. Box plot of per cent identity between BLASTN hits for *M. amalphitanum* integrase-related TE transcripts showed that high-copy hits represent MITEs (**Figure S14**). We hypothesize that actively proliferating TE copies can represent recent arrivals, possibly brought about by viruses or host-parasite interactions [54].

Our study provides a first view of the genomic content of one of the smallest insects currently known, the parasitoid wasp *M. amalphitanum*. In contrast to the expectation that the small body size, in combination with the parasitic lifestyle, should lead to significant reduction in the amount of genomic DNA and in gene content, we do not observe a drastic reduction in the overall genome size or in the number of expressed genes in comparison with larger parasitic wasps.

Of the rare cases of genes that were apparently lost from – or highly degenerated in – the *M. amalphitanum* genome, the most intriguing is *centrosomin (cnn)*, which is universally present in insects, including *T. pretiosum.* In *Drosophila melanogaster*, Cnn has important roles at the centrosome in mitotic spindle formation, cytoskeleton organization and neuronal morphogenesis [55, 56], although these functions may not be indispensable because this species (and possibly other insects) possesses centrosome-independent mechanisms for spindle nucleation [57]. A fungal homolog of Cnn, called *anucleate primary sterigmata*, is involved in nuclear migration [58–60]. We speculate that the loss of this gene is linked to the unique biological feature of widespread neuron denucleation in this tiny parasitoid wasp.

Surprisingly, transposable element dynamics over time was found to differ greatly between the analyzed species, with *M. amalphitanum* displaying a relatively recent dramatic decline in TE activity preceded by a burst, a pattern not observed in other parasitoid wasps. The decline in TE activity may have been associated with evolution of additional *Ago* and *Piwi* copies, not present in *T. pretiosum*, which could have reinforced the genome defense machinery to prevent uncontrolled TE expansion.

The relationship between body size and genome size has been discussed for a long time. Significant correlations of these values have been described for flatworms and copepods [16]; by contrast, such correlations were not found in ants [61]. Our results show that body size reduction in hymenopterans, while being associated with certain genomic adaptations, is not accompanied by greatly decreased transcriptomic and genomic complexity. This observation begs the question of how miniaturization is encoded genetically. We hypothesize that changes in regulatory sequences, rather than gene content, were important in the process of body size reduction, similar to mechanisms of morphological evolution that have driven adaptive diversification in all animals, great or small [62].

## Acknowledgements

This work has been carried out using computing resources of the federal collective usage center Complex for Simulation and Data Processing for Mega-science Facilities at NRC “Kurchatov Institute” (ministry subvention under agreement RFMEFI62117X0016), http://ckp.nrcki.ru/. Support of this project was provided by the Russian Scientific Foundation (RSF) grant #14-24-00175 and RSF grant #14-50-00060. B.L. was supported by the Rosenthal Brown-MBL internship and the REU supplement to NSF MCB-1121334 to I.A. R.B. was supported by a European Research Council Consolidator Grant (615094).

## Data availability

All genome and transcriptome assemblies and all SRA sequence data are publicly available at the NCBI BioProject: PRJNA344956.

## Competing interests

The authors declare that they have no competing interests.

