## Supplementary File for "A draft genome sequence of the miniature parasitoid wasp, *Megaphragma amalphitanum*"

### Supplementary Notes

#### A. Next-generation sequencing

##### A1. *Megaphragma amalphitanum* DNA extraction and DNA library preparation

We used *Megaphragma amalphitanum* (Hymenoptera: Trichogrammatidae) imago individuals (males and females) reared in the laboratory from eggs of *Heliothrips haemorrhoidalis* (Thysanoptera: Thripidae) collected in Santa Margherita (Northern Italy). The chitin exoskeleton of insect bodies was destroyed using an ultrasonic bath (Elma Ultrasonic S10, Germany) with an intensity of 37 kHz. DNA was extracted using NucleoSpin Tissue XS kit (Macherey-Nagel, Germany). DNA-libraries were constructed using Ovation Ultralow Systems V2 kit (NuGEN, USA) following the manufacturer's protocol with 13 cycles of library amplification (**Table S1**). Mate-pair libraries could not be constructed due to the limite.d amount of biological material

##### A2. *M. amalphitanum* RNA extraction and cDNA library preparation

Ten *M. amalphitanum* imagoes (males and females) were resuspended in TRIzol® (Thermo Fisher Scientific, USA) and extracted following the manufacturer's protocol. A steel homogenizer was used to quickly disrupt the chitin exoskeleton.

A total of ~10 ng *M. amalphitanum* RNA was extracted, and cDNA libraries were constructed using Ovation RNA-Seq System V2 kit (NuGEN, USA) with poly(A) enrichment following the manufacturer's protocol with 7 cycles of library amplification. The cDNA library size was approximately 330 bp.

The final library met all quality metrics as defined by Illumina, and library quantitation was performed on an Agilent 2100 Bioanalyzer with a High-Sensitivity DNA chip (Agilent Technologies, USA) prior to sequencing.

**A3. Genome and cDNA sequencing**

*M. amalphitanum* genome libraries were sequenced using 150-bp read length in a single flow cell on an Illumina Hiseq1500 (Illumina, USA). A total of 102,188,833 Illumina paired-end reads were generated and used for *de novo* genome assembly.

Additionally, genomic DNA-libraries from the thorax and abdomen of *M. amalphitanum* (SRR5982987) and from the heads of *M. amalphitanum* (SRR5982986) were prepared and sequenced. In total, 79,317,970 (paired-end sequencing: 2×100 bp) and 85,409,775 (single-end sequencing: 50 bp) DNA reads were sequenced and were used for *M. amalphitanum* coverage increase and as additional evidence during the search for missing (**Table S1**).

*M. amalphitanum* cDNA libraries were sequenced using 150-bp read length in a single flow cell on an Illumina Hiseq1500 (Illumina, USA). The Illumina sequencing generated a total of 59,790,973 paired-end reads, which were used for *de novo* transcriptome assembly.

**B. Data analysis**

**B1. Genome *de novo* assembly.**

The output from Illumina sequencing of the genomic DNA library (source format *.fastq) was used for *de novo* genome assembly. To remove contaminants, we used BBDuk software (v. 37.08), which is included in the BBMap package (v. 35.92), using the bacterial, fungal, plant, virus and “other” databases (http://jgi.doe.gov/data-and-tools/bbtools/bb-tools-user-guide/). Accession numbers of contaminating sequences are presented in Supplementary Dataset 1. To assemble the complete genome of *M. amalphitanum,* we used 102,188,833 paired-end reads. Genome assemblies have been constructed using different assembly algorithms (ABySS (v. 1.5.2)[1], Velvet (v. 1.2.10)[2], CLC genomic workbench software (version 5.5), SOAP (v. 2.0.1)[3] and SPAdes (v.3.6.1)[4]), and their performance was compared to each other (**Figure S1**).

At the first step of genome assembly, we merged the reads using Pear software (v. 0.9.5)[5]. As a result, up to 65% of the reads were merged, and genomic paired reads were quality trimmed and filtered. Before making a *de novo* assembly with the ABySS assembler, we used the Quake package (v. 0.3)[6], which fixes the sequencing errors. Then we launched ABySS with the k parameter varying from 40 to 92 in increments of 4. The best assembly was achieved for k = 80. The results are presented in **Table S2**.

For the SOAP aligner, we used the SOAPec_v2.01 pipeline for sequencing error correction. K-mer length estimation was done with the KmerGenie program [7], with the resulting k equal to 53. Predicted assembly size was 487,813,340 bp.

For SPAdes assembly (SPAdes v.3.6.1) (http://bioinf.spbau.ru/ru/SPAdes) we used the parameter "-careful", which minimizes the number of errors in the final contigs using BayesHammer error corrector and automatic k-mer estimation [8]. Also, we used a commercial *de novo* assembler in the CLC genomic workbench software package (version 5.5) with default parameters (**Table S2**).

For the final assembly, we used SPAdes. With the utility Data Preparation Module from SOAP, 94,687 scaffolds were built with N50 equal to 10,296 nucleotides, and the total assembly size of 346 million nucleotides (**Table S3**).

##### B2. *M. amalphitanum* and *Ceratosolen solmsi* transcriptome *de novo* assembly

Illumina RNA sequencing generated a total of 59,790,973 paired-end reads. For further analysis after filtration and contaminant removal, 50,529,351 paired-end reads were used for *M. amalphitanum*. To remove contaminants, we used BBDuk software (v. 37.08), which is included in the BBMap package (v. 35.92), using the bacterial, fungal, plant, virus and "other" databases (http://jgi.doe.gov/data-and-tools/bbtools/bb-tools-user-guide/). Accession numbers of contaminating sequences are presented in Supplementary Dataset 1.

Transcriptome *de novo* assembly was conducted using the Trinity software (v. 2.4.0)[9] with a default k-mer size. The Trinity software package combines three assembly algorithms: Inchworm, Chrysalis and Butterfly. Inchworm builds a K-mer dictionary from the reads, which leads to the construction of contigs. Chrysalis connects all overlapping contigs into components using the de Bruijn graph approach. In a final step, Butterfly simplifies all the generated graphs to report full-length transcripts and their alternatively spliced forms.

We assembled 46,841 contigs for *M. amalphitanum* and 62,786 contigs for the related parasitoid chalcid wasp *Ceratosolen solmsi*. These data were used in subsequent transcriptome annotation and radar plot construction using Excel. Radar plots (**Figures S7-S9**) demonstrate level of similarity between parasitoid wasps (*M. amalphitanum*, *C.*

*solmsi*, *D. alloeum*, *F. arisanus*, *C. vestalis*, *T. pretiosum)* based on gene expression. Assembly statistics is presented in **Table S5.**

**B3. *M. amalphitanum* and other parasitoid wasp transcriptome annotation**

Annotation of *M. amalphitanum, C. solmsi*, *Diachasma alloeum*, *Fopius arisanus*, *Cotesia vestalis*, and *Trichogramma pretiosum* transcriptome assemblies (**Table S6**) was performed using the Trinotate pipeline (https://trinotate.github.io/) from the Trinity package. All assembled contigs were searched against several databases (NCBI (Nr), Swissprot-Uniprot, Kyoto Encyclopedia of Genes and Genomes (KEGG), GO (Gene Ontology) and EggNog) using BLASTX[10] with an E-value cut-off set to 10−5. Gene open reading frames (ORFs) were predicted using Transdecoder (http://transdecoder.github.io/). We retained only predicted ORFs that were at least 100 amino acids long, either partial or complete. The remaining functional annotation was achieved using Trinotate. The Trinotate pipeline uses several software tools: Hmmer v.3.1b1, a protein domain identification (PFAM) software, Tmhmm v.2.0c prediction of transmembrane helices in proteins, Rnammer v.1.2 to predict ribosomal RNA, SignalP v.4.1 to predict signal peptide cleavage sites, prediction of Gene Ontology GOseq, and eggnog v.3.0 search for orthologous groups. Trinotate statistic is presented in **Table S7**.

The major gene GO categories (biological processes, molecular function, and cellular components) were selected for radar-plot construction in Microsoft Excel 2010 (**Figures S7-S9**).

#### B4. Missing genes and missing or rapidly evolving gene clusters in the *M. amalphitanum* genome

Genomic data (from SRA archive, see **Table S6**) of six parasitoid wasp species (*T. pretiosum*, *C. vestalis*, *Copidosoma floridanum*, *F. arisanus*, *Nasonia vitripennis*, *N. giraulti*) and *M. amalphitanum* DNA reads were mapped uniquely using bowtie2 onto the set of *Apis mellifera* annotated genes (PRJNA13343, PRJNA10625). We identified covered and uncovered genes, and compared the covered genes between these seven parasitoid wasp species from the Chalcidoidea and Ichneumonoidea superfamilies (unique reads were filtered by tag: *-xs*). Using this analysis, we found 115 genes that were present in larger parasitoid wasps but lost from *M. amalphitanum*. The number of genes that are present only in *M. amalphitanum* reached the saturation threshold during sequential addition of genomic data to the "non-miniaturized wasp panel" (2 to 6). At the same time, after addition of any other wasp data (instead of *M. amalphitanum* data) the number of unique genes went down to zero (**Figure S10**).

The comparison of variance analysis of nucleotide sequences for 78 homologs of missing genes of *M. amalphitanum* in nine randomly selected insects (*Bombus impatiens*, *Culex quinquefasciatus*, *Drosophila melanogaster*, *Lucilia cuprina*, *N. vitripennis*, *Solenopsis invicta*, *Mayetiola destructor*, *Megaselia scalaris*, *Atta cephalotes*) and the variance of amino-acid sequences of randomly selected 78 genes of *A. mellifera* and the same set of insects that was repeated 10 times shows that most of the *M. amalphitanum* missed genes apparently are not lost, but differ significantly (**Figure S11**). A t-test for homologs of missing genes and for the average bootstrapping sample of randomly selected genes of *A. mellifera* gave a *p-value* = 0.0007.

The reciprocal best BLAST hits for 115 genes were analyzed with the following parameters: maximum E-value threshold = 1e-5 and coverage of 30% of any of the protein sequences in the alignments as described in Ward with colleagues [11]. Only 78 homologs of genes that were not covered by *M. amalphitanum* reads were retained after this procedure. These genes were subjected to further scrutiny by retrieving a wasp ortholog and subsequently using its amino acid sequence as a query in a TBLASTN search of the *M. amalphitanum* genome database (E=1e-5). In this way, we could identify the presence of highly fragmented genes and/or those genes that are subject to rapid evolution.

OrthoMCL [12] with parameters recommended in previous studies [13] was used for identification of missing or rapidly evolving gene clusters in the *M. amalphitanum* and other Chalcidoidea genomes.

**B5. Search for chemoreceptor genes in *M. amalphitanum***

All of the 94,687 scaffolds were used in searches for chemoreceptor genes and for comparison with orthologous genes from other insect genomes in the NCBI (National Center for Biotechnology Information, USA) database. We used BLASTX with parameter -minIdentity = 30 and -minScore = 25 to search for Ionotropic Receptors (IRs), Odorant Receptors (ORs) and Gustatory Receptors (GRs). The list of accession numbers of chemoreceptor genes used as queries is presented in Supplementary Dataset 2.

**B6. Transposable element (TE) *de novo* identification and analysis**

For *de novo* TE library construction, we used the REPET package [14] which combines three mutually complementing repeat identification tools (RECON,

GROUPER and PILER), yielding a combined repeat library with the average consensus sequence length of 1.66 kb (in the range 157-14,640 bp). The libraries were subject to additional classification with the RepeatClassifier tool from the RepeatMasker package (www.repeatmasker.org) using the Repbase database of eukaryotic repetitive elements (www.girinst.org) [15]. RepeatMasker was also used to build the corresponding TE landscape divergence plots. The calculated Kimura 2-parameter distances were additionally subjected to CpG correction because of the presence of a functional DNA methylation system in all analyzed hymenopterans (both *Dnmt1* and *Dnmt3* are present in *M. amalphitanum*). Note that while our TE content estimates in *N. vitripennis* and *Polistes dominula* using *de novo* repeat libraries were very close to the published analyses [16, 17], the previously reported TE content in *Polistes canadensis* represents a substantial underestimate, since it included only RepBase entries from Apocrita [18].

Due to the higher-than-usual proportion of "unknown" sequences in the classified *M. amalphitanum* repeat library (11%), we extracted the "unknown" sequences and subjected then to re-classification. This resulted in removal of a few multicopy host genes and re-assignment of known TEs which were not recognized by RepeatClassifier (e.g. MITEs). In the end, the "unknown" repeat content was reduced to an acceptable number of 7% (**Figure S12**).

Transcriptionally active TEs were extracted from GO matches corresponding to DNA integration, and scanned against Repbase (http://www.girinst.org/censor/) [15], which contains curated full-length TE consensus sequences from multiple species, including six hymenopteran insects. Fragments were grouped according to their best database matches for assignment to different families, with 10-20 fragmented contigs spanning each family across the entire length. This procedure also permitted detection of recently arrived TEs present in less than 3 copies per genome, which is the limit of detection for *de novo* genomic repeat library construction. Copy numbers and the degree of divergence of transcribed copies were estimated from BLASTN searches of transcripts against genomic DNA (-evalue 10e-5).

TE defense genes: To identify candidate genes in the TE silencing system in the genomes of *M. amalphitanum* and *T. pretiosum*, we created a database of genes from NCBI reference sequences of Piwi-Argonaute and Dicers proteins. This dataset was used in reciprocal BLAST searches against the assemblies of *M. amalphitanum* and *T. pretiosum* (using TBLASTN). Initial BLAST outputs were curated, merging HSP and annotating gene boundaries to identify homologs. Each potential homolog was then compared to the NCBI nr database (BLASTX) for final confirmation. Selected sequences were run against the SNAP *ab inito* gene prediction software [19] for identification of open reading frames. NCBI's Conserved Domain Database was launched for conserved domain annotation of predicted proteins.

Multiple alignments of CDS sequences were performed using Muscle v3.8 [20] with default settings. Phylogenetic trees were generated under the maximum likelihood criterion using PhyML 3.0 (GTR model, NNI topological moves and likelihood branch supports) [21]. All manipulations of phylogenetic trees were performed using FigTree [22].

##### B7. Search for venom components in the *M. amalphitanum* transcriptomic data

The presence of homologues of *N. vitripennis* poison constituents in *M. amalphitanum* and other parasitoid wasps (*Megastigmus spermotrophus*, *N. vitripennis*, *C. solmsi*, *T. pretiosum*), was conducted using previously published venom data[23, 24] and the transcriptomes of chalcid wasps (**Table S6**). Each *N. vitripennis* venom protein query was compared with the four sets of transcriptome data using TBLASTN, with an E-value cut-off of 1e-07.

#### Supplementary Figures

**Figure S1.** *M. amalphitanum* genome assembly statistics using ABySS, SPAdes, CLC Genomics Workbench and Velvet software. K-mer sizes were matched for ABySS, SPAdes and Velvet; Note: CLC Genomics Workbench does not use k-mer size; CLC assembly was performed with default settings.

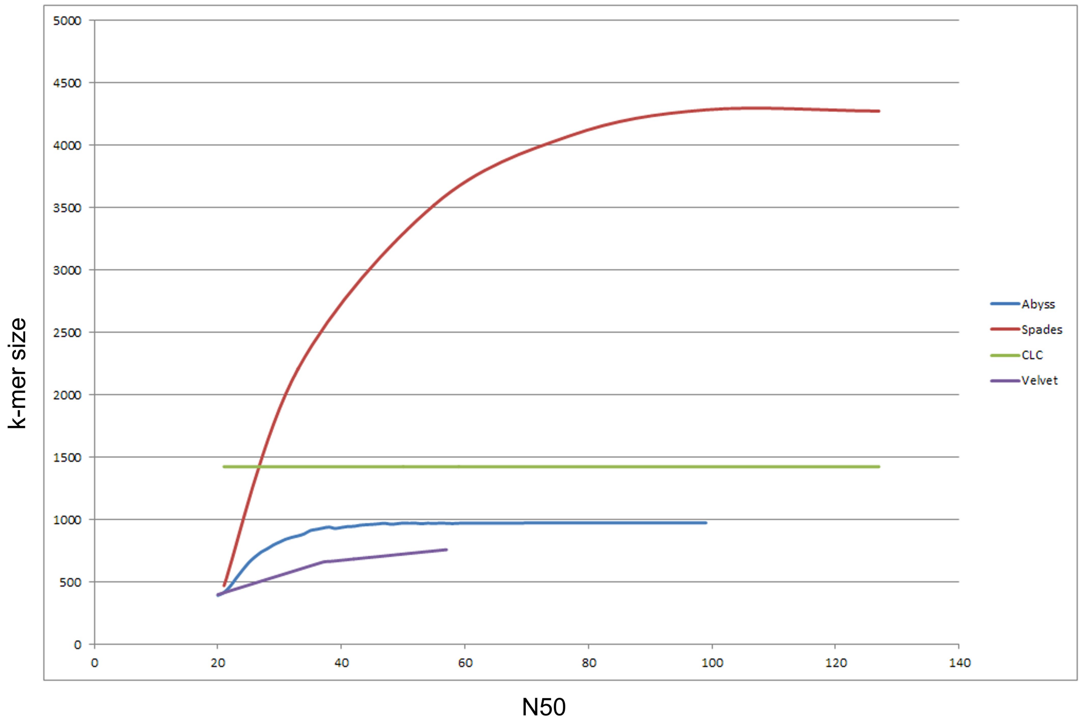

**Figure S2**. Gene ontology analysis of *M. amalphitanum* transcriptome for contigs with assigned GO: Biological processes.

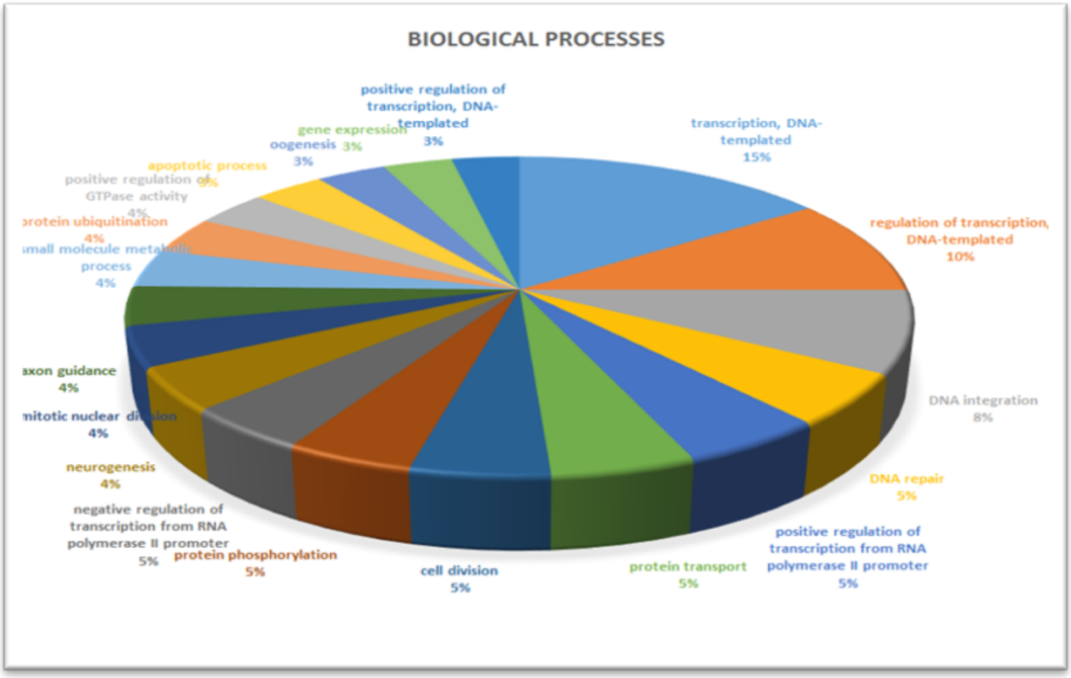

**Figure S3**. Gene ontology analysis of *M. amalphitanum* transcriptome for contigs with assigned GO: Molecular function.

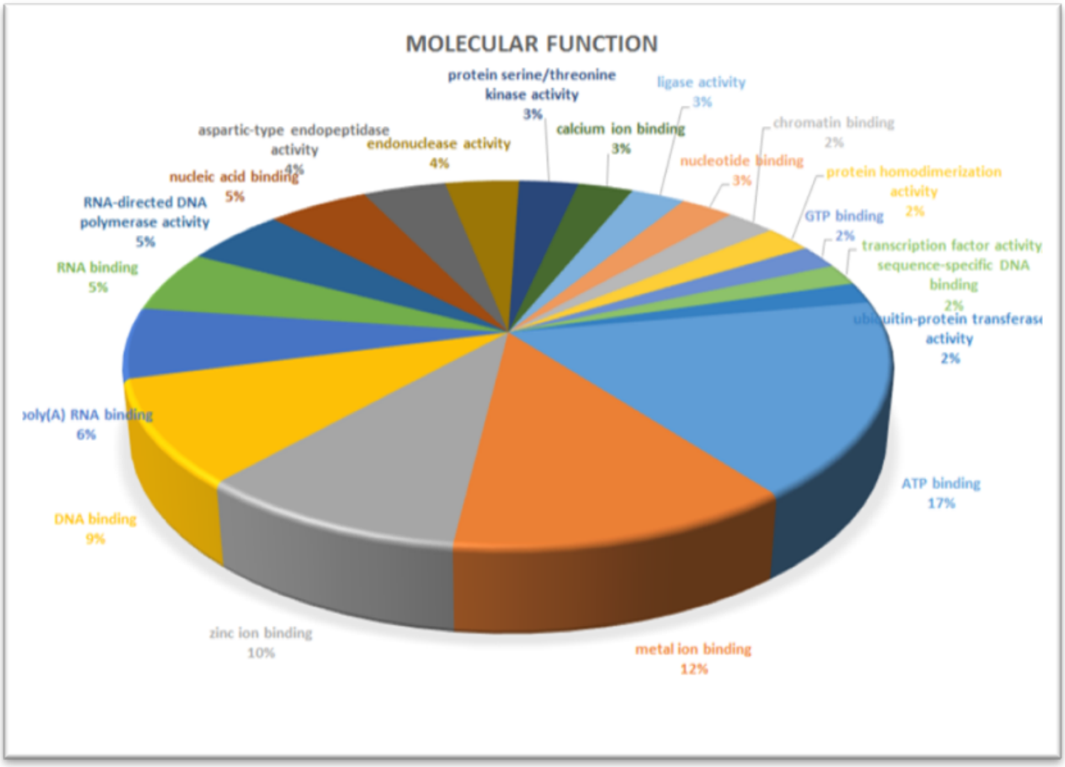

**Figure S4**. Gene ontology analysis of *M. amalphitanum* transcriptome for contigs with assigned GO: Cellular components.

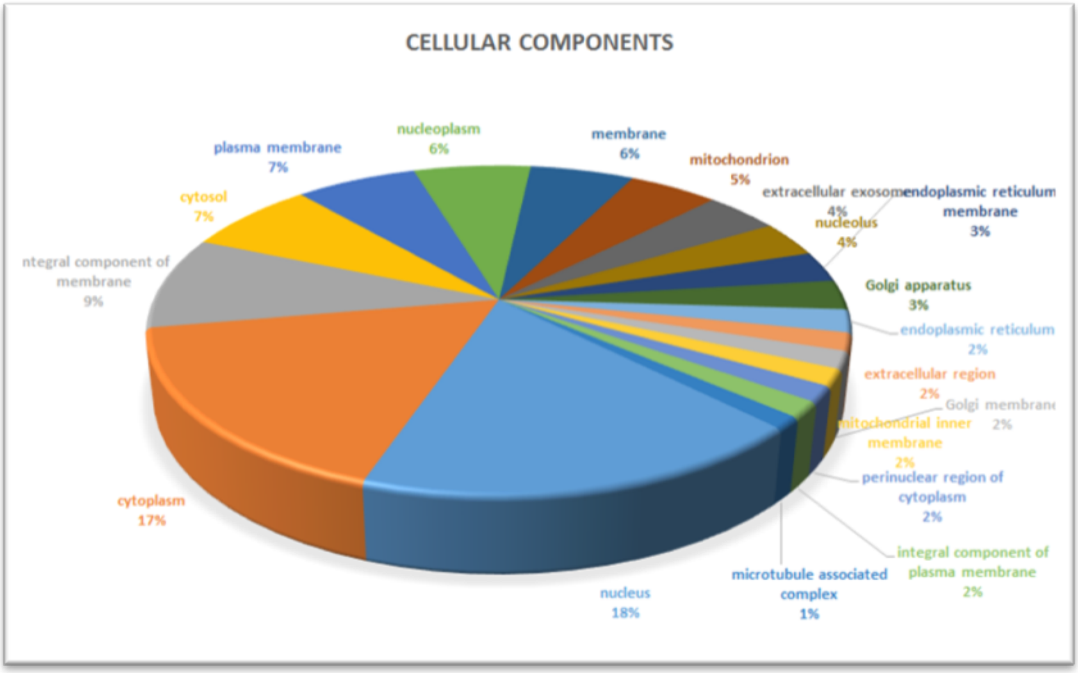

**Figure S5**. The Clusters of Orthologous Groups (COG) for *M. amalphitanum* transcriptome (top pathways).

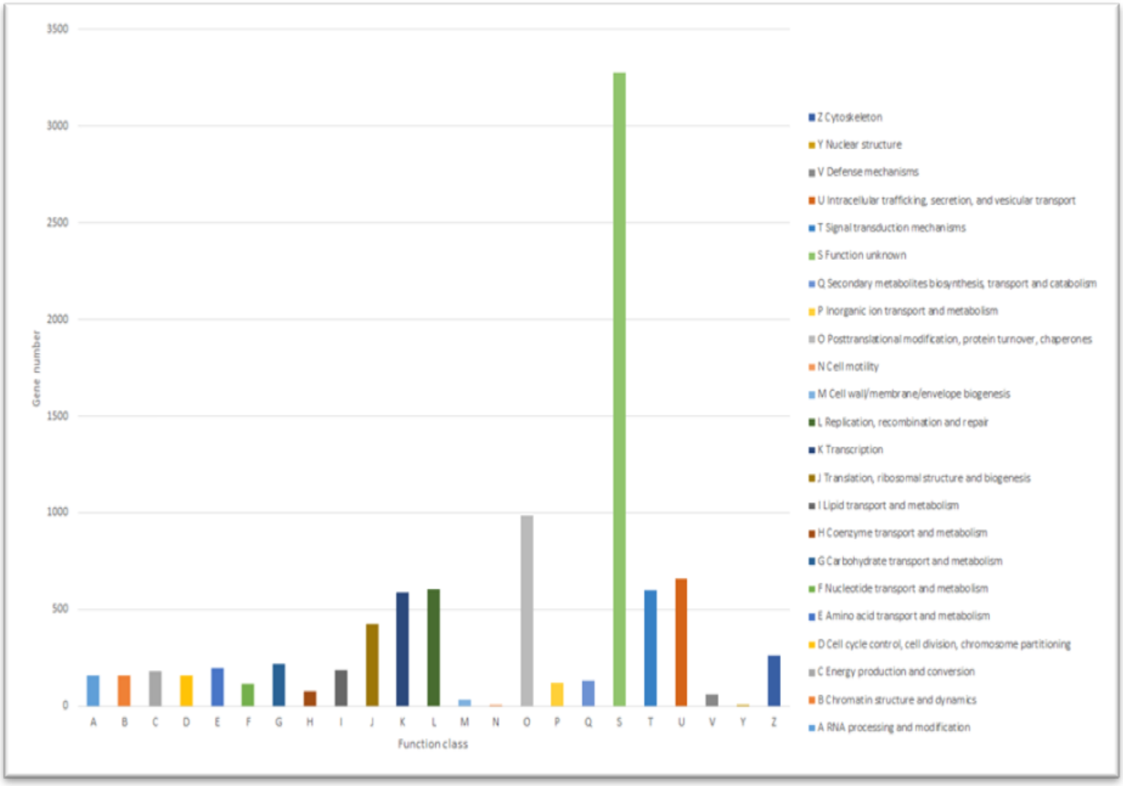

**Figure S6.** KEGG pathway analysis for the *M. amalphitanum* transcriptome.

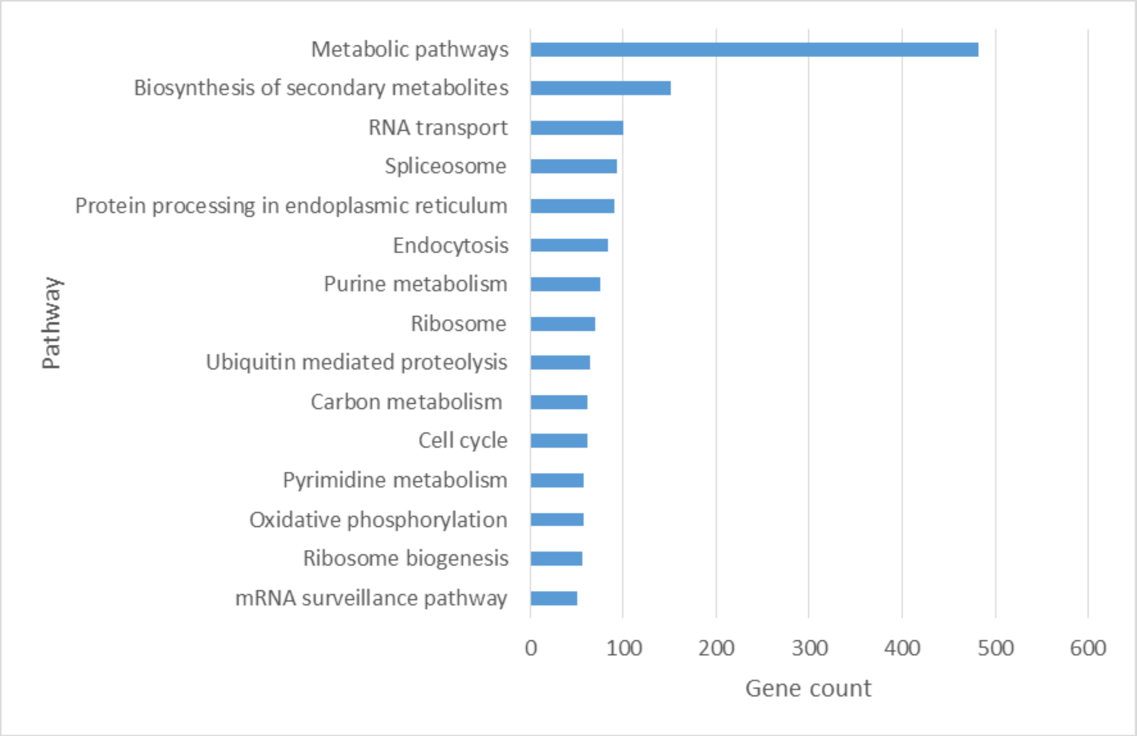

**Figure S7.** Radar plot for the *M. amalphitanum, C. solmsi*, *D. alloeum*, *F. arisanus*, *C. vestalis*, *T. pretiosum* transcriptome GO-category related to biological processes showing numbers of transcripts in this GO-category for six Chalcidoid species.

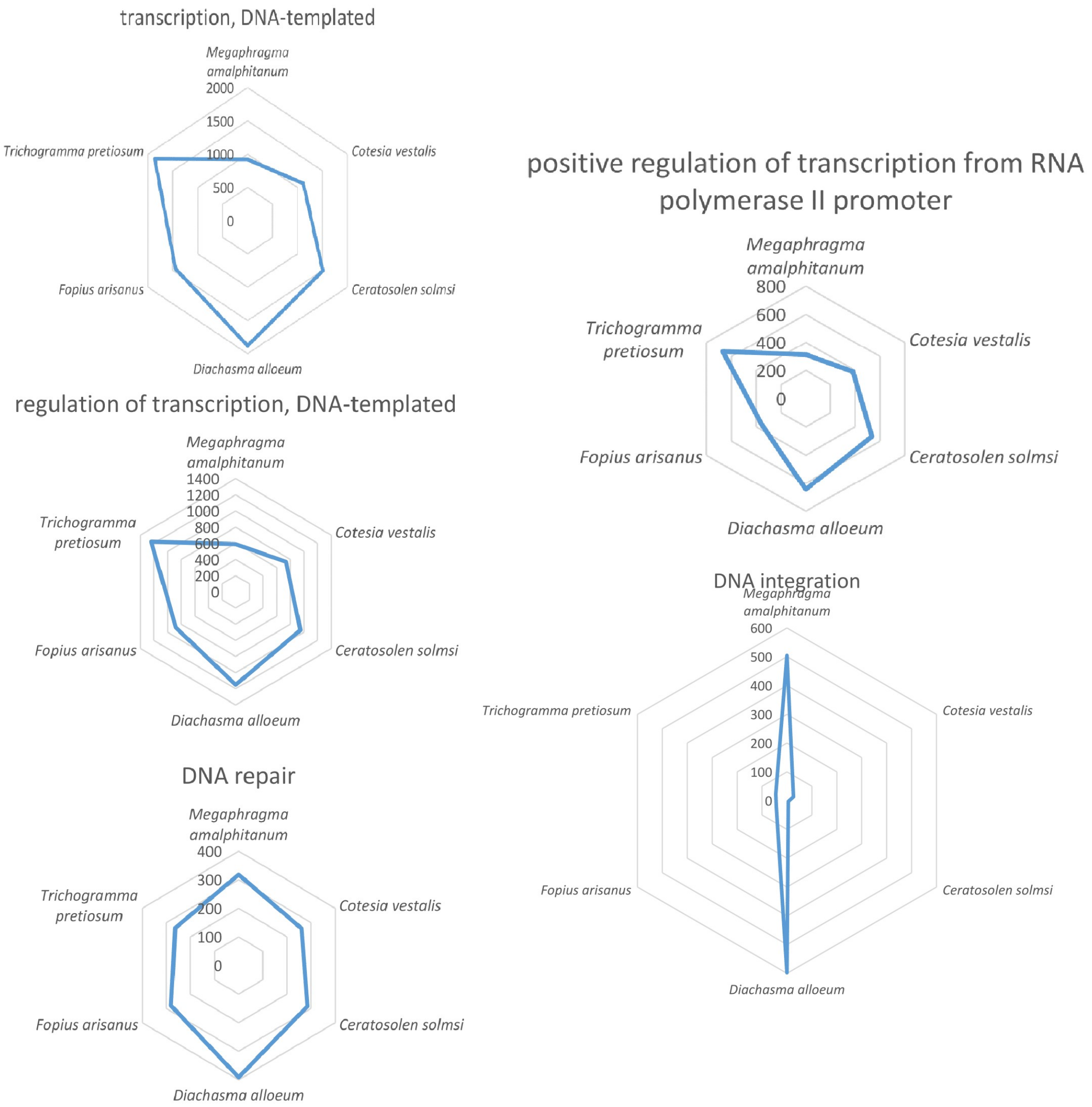

**Figure S8.** Radar plot for the *M. amalphitanum, C. solmsi*, *D. alloeum*, *F. arisanus*, *C. vestalis*, *T. pretiosum* transcriptome GO-category related to cellular components showing numbers of transcripts in this GO-category for six Chalcidoid species.

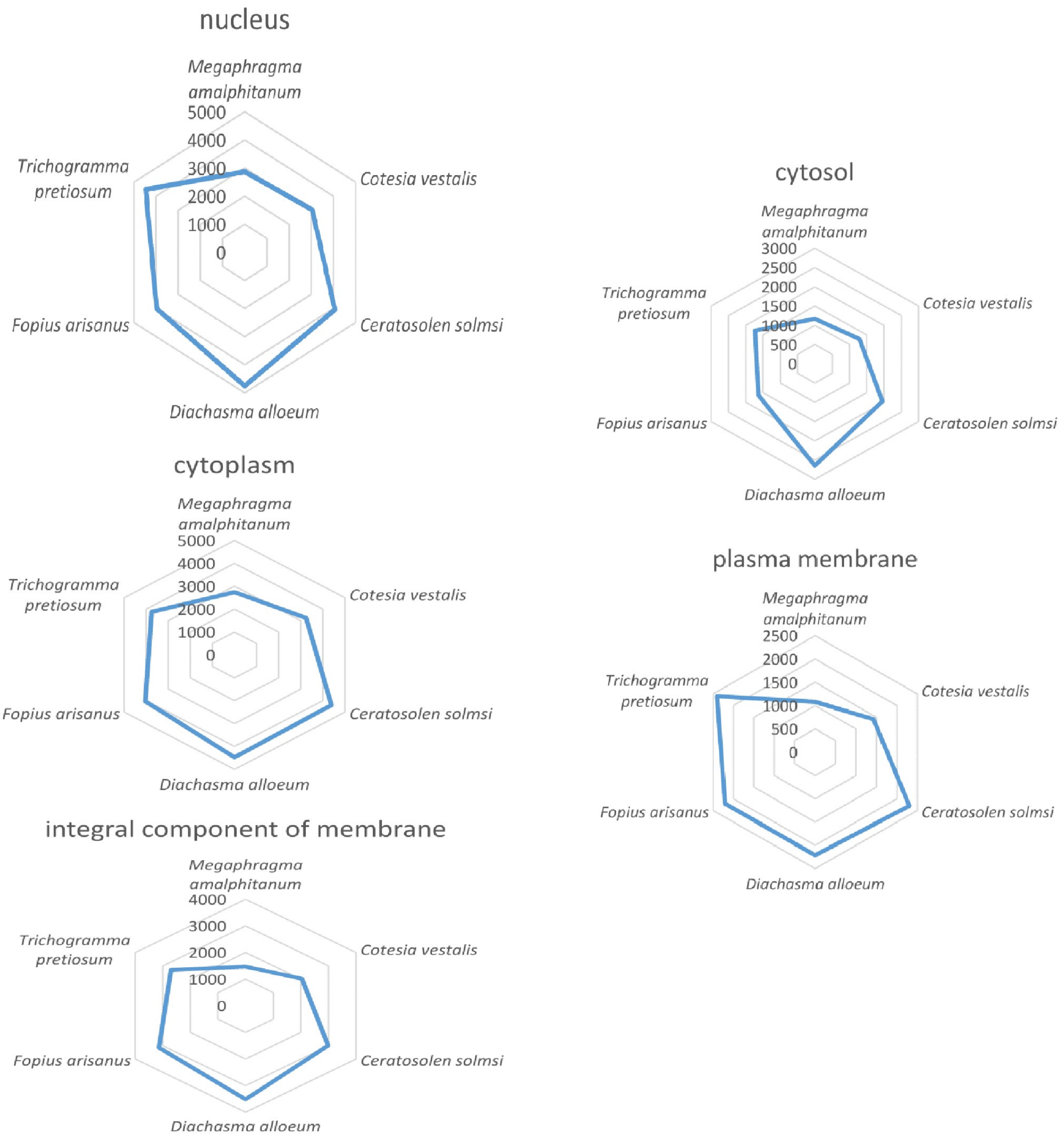

**Figure S9.** Radar plot for the *M. amalphitanum, C. solmsi*, *D. alloeum*, *F. arisanus*, *C. vestalis*, *T. pretiosum* transcriptome GO-category related to molecular processes showing numbers of transcripts in this GO-category for six Chalcidoid species.

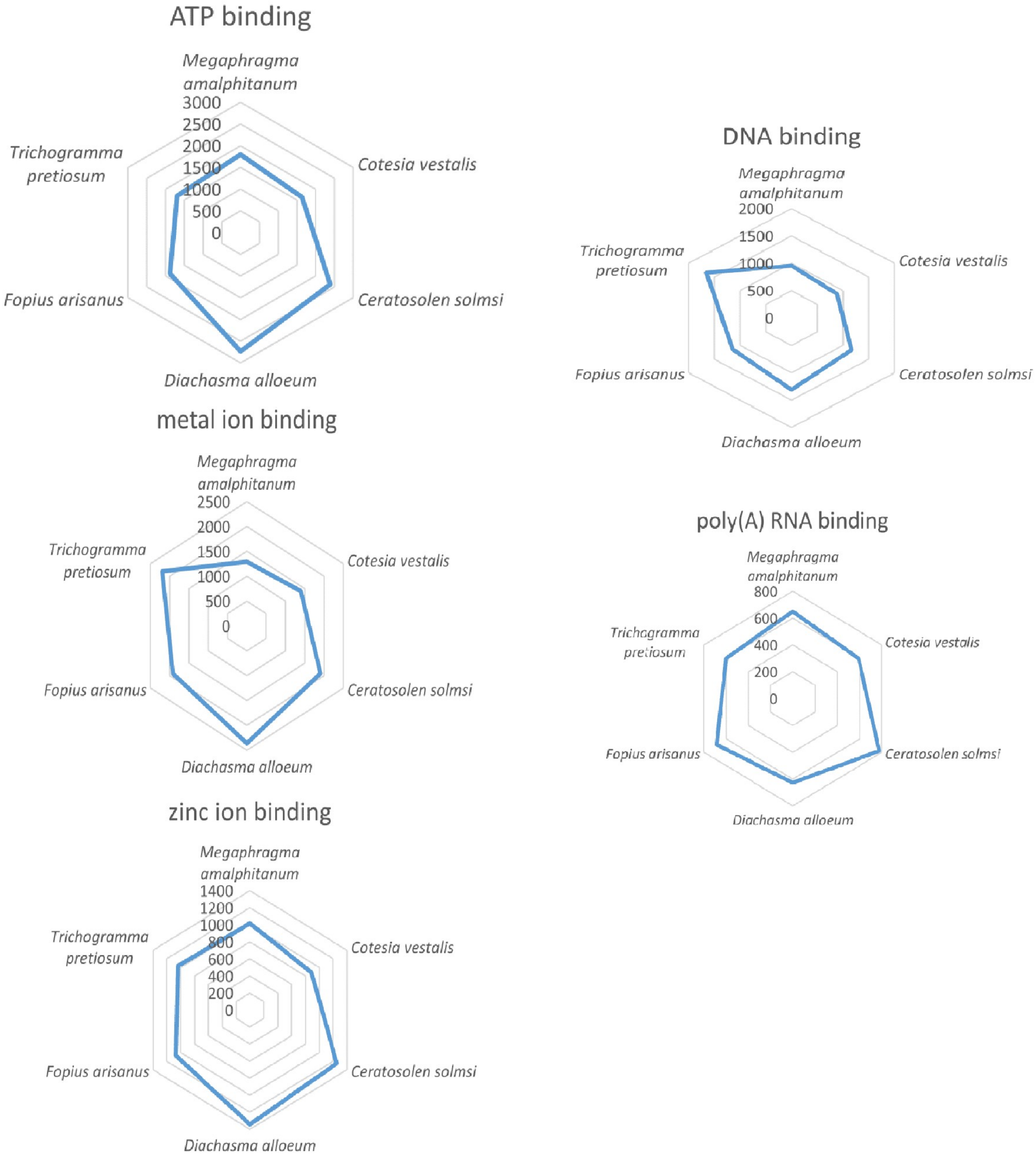

**Figure S10.** Missing genes in the *M. amalphitanum* genome. Y-axis: number of genes; X-axis: number of hymenopteran genomes analysed.

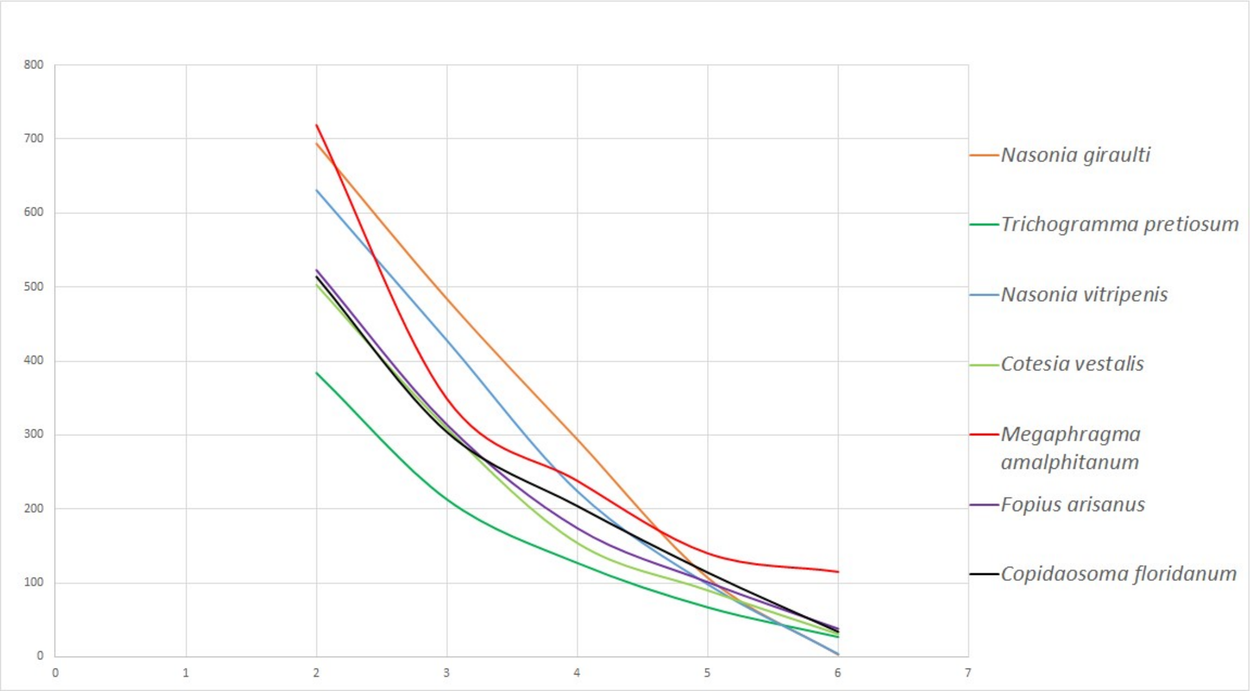

**Figure S11.** The comparison of variance analysis of the nucleotide sequences for 78 homologs of *M. amalphitanum* missing genes in nine randomly selected insects (*B. impatiens*, *C. quinquefasciatus*, *D. melanogaster*, *L. cuprina*, *N. vitripennis*, *S. invicta*, *M. destructor*, *M. scalaris*, *A. cephalotes*) (black line) and the variance of the nucleotide sequence of randomly selected 78 genes of *A. mellifera* and the same set of insects (red lines) that was repeated 10 times.

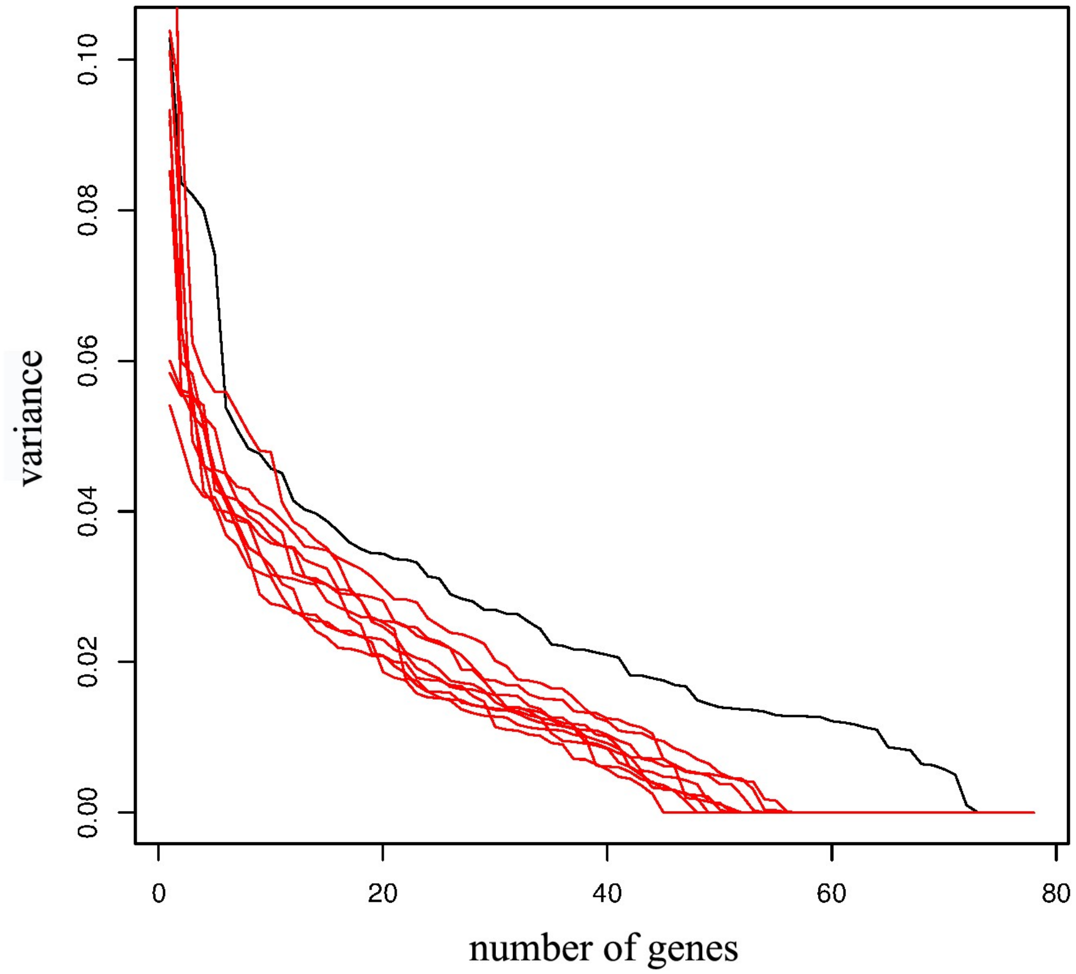

**Figure S12.** Effects of re-classification of “unknown” repeats in the *de novo* library for *M. amalphitanum* and *P. dominula* (**Supplementary Notes B6**). v2, re-classified.

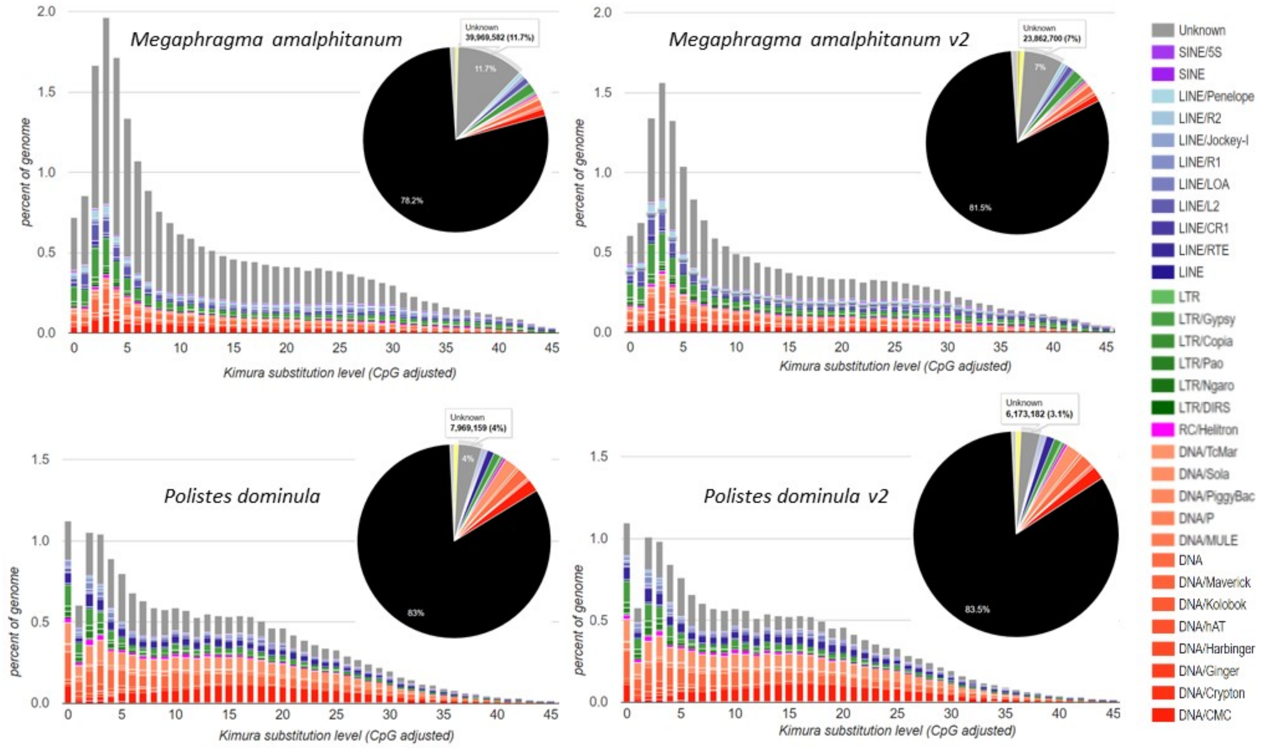

**Figure S13.** Maximum likelihood analysis of phylogenetic relationships among eukaryotic Dicer homologs from animals, plants, and fungi. *M. amalphitanum* and *T. pretiosum* Dcr-1 and Dcr-2 homologs are denoted by red dots. Multiple alignments of CDS sequences were performed using Muscle v3.8 [20] with default settings. Phylogenetic trees were generated under the maximum likelihood criterion using PhyML 3.0 (GTR model, NNI topological moves and likelihood branch supports) [21]. All manipulations of phylogenetic trees were performed using FigTree [22]. Scale bar, nucleotide substitutions per site

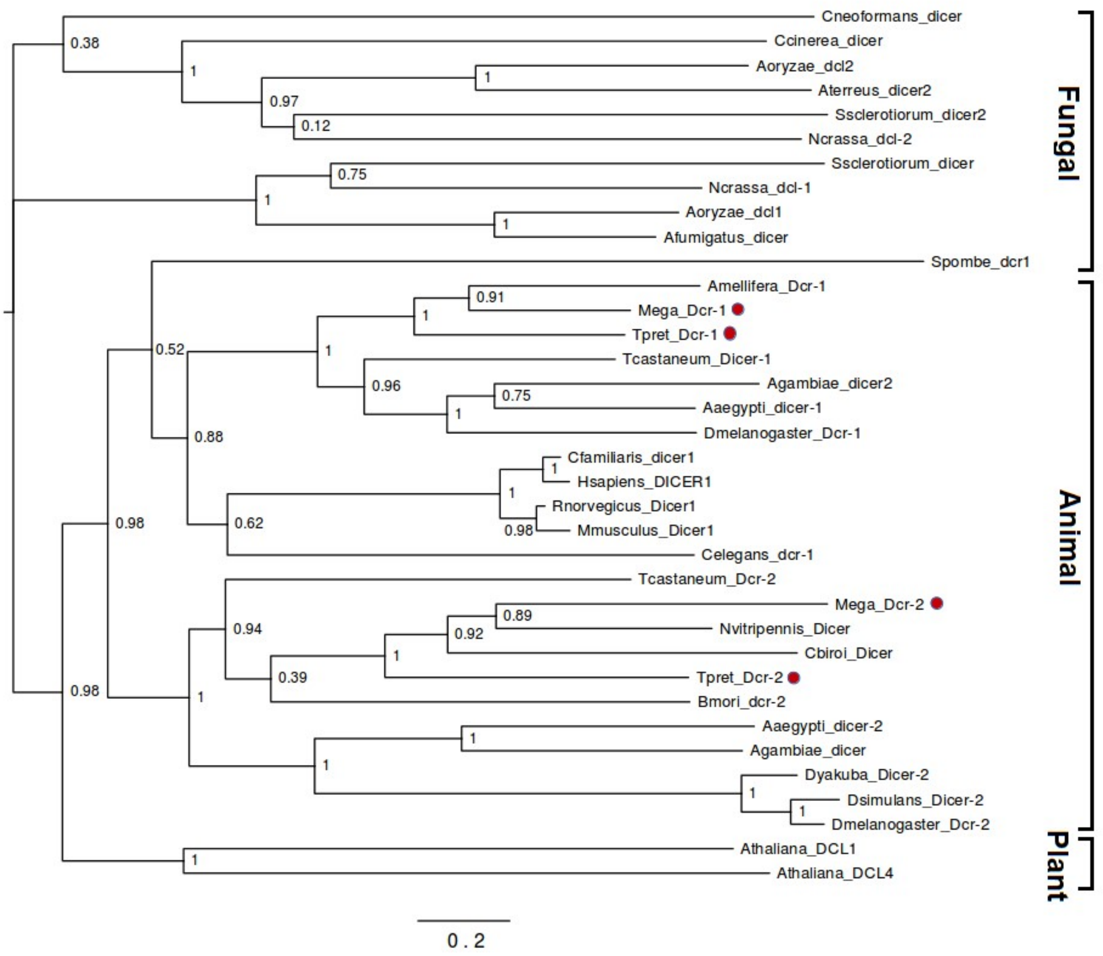

**Figure S14.** Box plot of percent identity between BLASTN hits for *M. amalphitanum* integrase-related TE transcripts, binned by copy count. High-copy hits represent MITEs.

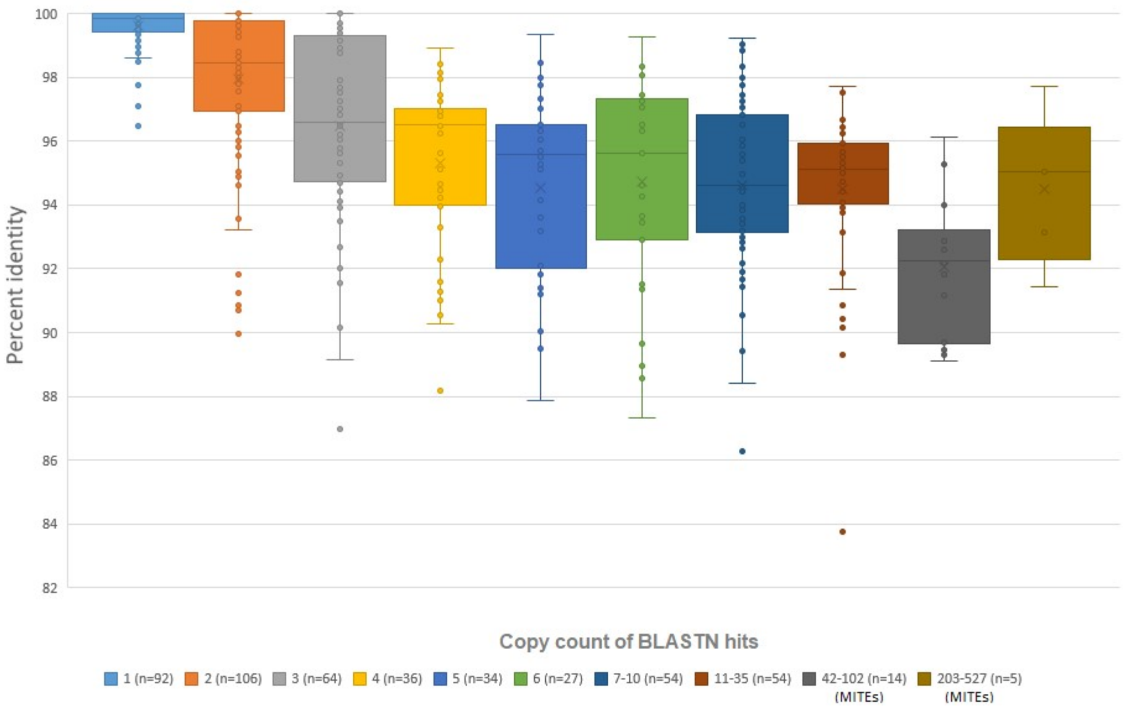

**Figure S15.** An overview of the missing gene analysis pipeline and its results

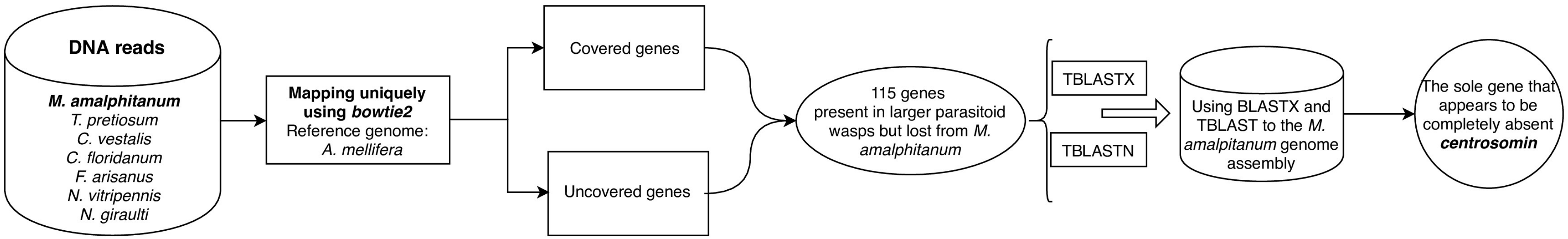

#### Supplementary Tables

**Table S1**. Paired-end DNA-libraries used for *M. amalphitanum* genome sequencing.

| Library name | Library concentration, Qubit, ng/µl | Average library size, Agilent 2100 Bioanalyzer with a High-Sensitivity DNA chip | SRA accession |
|---|---|---|---|
| DNA-library1 – whole insect (ten individuals) | 1.84 | 350 bp | SRR4340083 |
| DNA-library2 – body: thorax and abdomen (ten individuals) | 11.3 | 315 bp | SRR5982987 |
| DNA-library3 – head (ten individuals) | 1.8 | 334 bp | SRR5982986 |

**Table S2.** *M. amalphitanum* genome assembly statistics using ABySS, SPAdes, CLC and Velvet software (contigs).

| n | n:N50 | N50 | Maximum contig length, bp | Summary assembly size, bp | *De novo* assembler |
|---|---|---|---|---|---|
| 541950 | 31780 | 869 | 19313 | 2.91E+08 | Velvet |
| 484255 | 53248 | 1425 | 33321 | 3.07E+08 | CLC |
| 553157 | 20847 | 4285 | 56202 | 3.58E+08 | SPAdes |
| 4.58E+06 | 60283 | 974 | 21336 | 2.20E+08 | ABySS |

**Table S3.** *M. amalphitanum* genome final assembly statistics (scaffolds).

| n | n:N50 | N50 | Maximum contig length, bp | Cumulative assembly size, bp |
|---|---|---|---|---|
| 94687 | 7843 | 10296 | 895906 | $3.46\times10^{8}$ |

**Table S4.** Evaluation of the *M. amalphitanum* genome and transcriptome assemblies using the BUSCO (benchmarking universal single-copy orthologs) arthropod gene set.

| *M. amalphitanum* | Complete (%) | Duplicated (%) | Fragment (%) | Missing (%) |
|---|---|---|---|---|
| Genome | 64.74% | 4.37% | 27.70% | 7.55% |
| Transcriptome | 24.15% | 5.19% | 29.23% | 41.42% |

**Table S5.** *M. amalphitanum* and *C. solmsi* transcriptome assembly statistics using Trinity software (contigs).

| N | n:N50 | N50 | Maximum contig length, bp | Summary assembly size, bp | Wasp species |
|---|---|---|---|---|---|
| 46841 | 13109 | 633 | 9503 | $3.74\times10^{7}$ | *M. amilphitanum* |
| 62786 | 12699 | 724 | 15263 | $3.64\times10^{7}$ | *C. solmsi* |

**Table S6.** Reference data sets used for *M. amalphitanum* genome and transcriptome data analysis.

| Parasitoid wasp species used in analysis | Taxonomy (Suborder, Superfamily, Family) | Body size, mm | Genome size, Mbp | Neuron number in the brain | **Source link** | **Data type** | **Usage** |
|---|---|---|---|---|---|---|---|
| *Megaphragma amalphitanum* | Apocrita; Chalcidoidea; Trichogrammatidae | 0.25 | 346 | 4600 | https://www.ncbi.nlm.nih.gov/bioproject/PRJNA344956 | SRA reads | Transcriptome and Genome Data Analysis |
| *Trichogramma pretiosum* | Apocrita; Chalcidoidea; Trichogrammatidae | 0.5 | 195.1 | 18000 | https://www.ncbi.nlm.nih.gov/bioproject/275661 | SRA reads | Transcriptome Data Analysis |
| *Ceratosolen solmsi* | Apocrita; Chalcidoidea; Agaonidae | 2.7 | 278 | No data | http://sra.dnanexus.com/studies/SRP029703/experiments | SRA reads | Transcriptome Data Analysis: 62,786 contigs assembled |
| *Copidosoma floridanum* | Apocrita; Chalcidoidea; Encyrtidae | 1.2 | 555 | No data | https://www.ncbi.nlm.nih.gov/sra/SRR947009 https://www.ncbi.nlm.nih.gov/sra/SRR947010 | SRA reads | Genome Data Analysis |
| *Nasonia vitripennis* | Apocrita; Chalcidoidea; Pteromalidae | 2.2 | 295.8 | No data | https://www.ncbi.nlm.nih.gov/assembly/GCF_000002325.3 | Genome | Genome Data Analysis |
| *Nasonia giraulti* | Apocrita; Chalcidoidea; Pteromalidae | 2.3 | 283.6 | No data | https://www.ncbi.nlm.nih.gov/assembly/GCA_000004775.1 | Genome | Genome Data Analysis |
| *Diachasma alloeum* | Apocrita; Ichneumonoidea; | 4.2 | 388.7 | No data | https://www.ncbi.nlm.nih.gov/nuccore/GECN0 | Complete transcriptome | Transcriptome Data Analysis: |

| | | | | | | | |
|---|---|---|---|---|---|---|---|
| | Braconidae | | | | 0000000.1 | | 131,607 contigs; 112,635,971 bp total |
| *Fopius arisanus* | Apocrita; Ichneumonoidea; Braconidae | 4.5 | 153,6 | No data | https://www.ncbi.nlm.nih.gov/sra/SRR1560675 https://www.ncbi.nlm.nih.gov/sra/SRR1560673 https://www.ncbi.nlm.nih.gov/sra/SRR1560668 https://www.ncbi.nlm.nih.gov/nuccore/748430103 | Complete transcriptome and SRA reads | Genome Data Analysis. Transcriptome Data Analysis15,346 contigs; 50,620,881 bp total |
| *Cotesia vestalis* | Apocrita; Ichneumonoidea; Braconidae | 1.9 | 131,9 | No data | https://www.ncbi.nlm.nih.gov/sra/SRR2029645 https://www.ncbi.nlm.nih.gov/nuccore/511518236 | Complete transcriptome | Genome Data Analysis. Transcriptome Data Analysis: 30,024 contigs; 27,114,579 bp total; teratocyte (extraembryonic cell) |
| *Megastigmus spermotrophus* | Apocrita; Chalcidoidea; Torymidae | 2.8 | No data | No data | https://www.ncbi.nlm.nih.gov//bioproject/PRJNA274192 | SRA reads | Transcriptome Data Analysis |
| Reference Hymenoptera | - | - | - | - | | | |

| | | | | | | | |
|---|---|---|---|---|---|---|---|
| species | | | | | | | |
| *Apis mellifera* | Apocrita; Apoidea; Apidae | 15 | 246.9 | 850000 – 1200000 | http://metazoa.ensembl.org/Apis_mellifera/Info/Index | Complete genome | Genome and transcriptome analysis |

**Table S7.** Trinotate statistics for *M. amalphitanum, C. solmsi*, *D. alloeum*, *F. arisanus*, *C. vestalis*, *T. pretiosum* transcriptome assemblies.

| Parasitoid wasp species | Number of transcripts used for annotation | Number of transcripts annotated by BLASTX | Gene ontology for BLASTX data | Number of transcripts annotated by EggNog database | Number of transcripts annotated by KEGG database | Number of transcripts annotated by BLASTP | Number of transcripts annotated by Pfam | Gene ontology for Pfam | Prediction of transmembrane helices in proteins (TmHMM) |
|---|---|---|---|---|---|---|---|---|---|
| *C. solmsi* | 63783 | 17816 | 17046 | 14833 | 14499 | 12592 | 10826 | 7136 | 2159 |
| *D. alloeum* | 135999 | 42492 | 39607 | 27183 | 33384 | 29569 | 28427 | 19335 | 6840 |
| *F. arisanus* | 22452 | 19290 | 16792 | 14685 | 14622 | 16843 | 16379 | 16792 | 4509 |
| *C. vestalis* | 31395 | 12693 | 11994 | 10537 | 10383 | 10578 | 9768 | 6421 | 2074 |
| *T. pretiosum* | 20818 | 17351 | 15802 | 13736 | 13609 | 15891 | 16040 | 11801 | 4308 |
| *M. amalphitanum* | 46841 | 12238 | 10721 | 8810 | 6130 | 8193 | 6808 | 4196 | 1197 |

**Table S8.** A set of 78 genes (paralogs and homologs) not covered by *M. amalphitanum* reads.

| *A. melifera* gene ID | *Drosophila melanogaster* gene ID | *D. melanogaster* homologous gene |
|---|---|---|
| GB45679 | FBgn0261823 | Additional sex combs [Source:FlyBase;Acc:FBgn0261823] |
| GB47271 | FBgn0022710 | Adenylyl cyclase 35C [Source:FlyBase;Acc:FBgn0022710] |
| GB53936 | FBgn0000228 | Blastoderm-specific gene 25D [Source:FlyBase;Acc:FBgn0000228] |
| GB42838 | FBgn0031883 | Caper [Source:FlyBase;Acc:FBgn0031883] |
| GB45937 | FBgn0013765 | centrosomin [Source:FlyBase;Acc:FBgn0013765] |
| GB44187 | FBgn0028387 | chameau [Source:FlyBase;Acc:FBgn0028387] |
| GB44210 | FBgn0037240 | Contactin [Source:FlyBase;Acc:FBgn0037240] |
| GB41346 | FBgn0041342 | CTP:phosphocholine cytidylyltransferase 1 [Source:FlyBase;Acc:FBgn0041342] |
| GB41346 | FBgn0035231 | CTP:phosphocholine cytidylyltransferase 2 [Source:FlyBase;Acc:FBgn0035231] |
| GB46429 | FBgn0000527 | ebony [Source:FlyBase;Acc:FBgn0000527] |
| GB50237 | FBgn0033354 | Fanconi anemia complementation group I homologue [Source:FlyBase;Acc:FBgn0033354] |
| GB45272 | FBgn0001987 | Gliotactin [Source:FlyBase;Acc:FBgn0001987] |
| GB55822 | FBgn0266136 | Guanylyl cyclase at 76C [Source:FlyBase;Acc:FBgn0266136] |
| GB42147 | FBgn0030600 | highwire [Source:FlyBase;Acc:FBgn0030600] |
| GB44850 | FBgn0005654 | latheo [Source:FlyBase;Acc:FBgn0005654] |
| GB43279 | FBgn0034282 | Mapmodulin [Source:FlyBase;Acc:FBgn0034282] |
| GB43420 | FBgn0261109 | marionette [Source:FlyBase;Acc:FBgn0261109] |
| GB52997 | FBgn0265988 | mauve [Source:FlyBase;Acc:FBgn0265988] |
| GB44910 | FBgn0027497 | MLF1-adaptor molecule [Source:FlyBase;Acc:FBgn0027497] |
| GB55787 | FBgn0002878 | mutagen-sensitive 101 [Source:FlyBase;Acc:FBgn0002878] |
| GB49559 | FBgn0016919 | no mechanoreceptor potential B [Source:FlyBase;Acc:FBgn0016919] |

| | | |
|---|---|---|
| GB43945 | FBgn0061200 | Nucleoporin 153kD [Source:FlyBase;Acc:FBgn0061200] |
| GB43591 | FBgn0026058 | Ods-site homeobox [Source:FlyBase;Acc:FBgn0026058] |
| GB41452 | FBgn0023517 | Phosphoglycerate mutase 5 [Source:FlyBase;Acc:FBgn0023517] |
| GB41452 | FBgn0035004 | Phosphoglycerate mutase 5-2 [Source:FlyBase;Acc:FBgn0035004] |
| GB46511 | FBgn0035405 | piefke [Source:FlyBase;Acc:FBgn0035405] |
| GB46270 | FBgn0025740 | Plexin B [Source:FlyBase;Acc:FBgn0025740] |
| GB54796 | FBgn0025334 | Putative homeodomain protein [Source:FlyBase;Acc:FBgn0025334] |
| GB41985 | FBgn0024941 | Regulator of G-protein signalling 7 [Source:FlyBase;Acc:FBgn0024941] |
| GB53284 | FBgn0011829 | Ret oncogene [Source:FlyBase;Acc:FBgn0011829] |
| GB42110 | FBgn0264087 | Slowpoke binding protein [Source:FlyBase;Acc:FBgn0264087] |
| GB44534 | FBgn0039141 | spastin [Source:FlyBase;Acc:FBgn0039141] |
| GB43591 | FBgn0024184 | unc-4 [Source:FlyBase;Acc:FBgn0024184] |
| GB40007 | | |
| GB40447 | | |
| GB40540 | FBgn0033916 | |
| GB41035 | FBgn0050421 | |
| GB41249 | | |
| GB41330 | | 26S proteasome complex subunit DSS1 |
| GB41486 | FBgn0035421 | |
| GB42383 | | |
| GB44289 | | |
| GB44766 | | |
| GB45063 | | LIM/homeobox Lhx9-like |
| GB45314 | | |
| GB45456 | | |
| GB45501 | | |
| GB46013 | FBgn0032010 | Mucin-1-like/Nucleoporin NSP1-like |
| GB46267 | | |

| | | |
|---|---|---|
| GB46273 | | |
| GB46668 | | |
| GB46705 | | |
| GB46908 | FBgn0261550 | |
| GB47425 | FBgn0037525 | |
| GB47871 | | |
| GB47927 | | |
| GB48315 | | |
| GB48563 | FBgn0037094 | |
| GB48783 | | |
| GB48948 | FBgn0032648 | |
| GB48948 | FBgn0032647 | |
| GB48948 | FBgn0032649 | |
| GB49002 | | |
| GB49164 | | |
| GB49318 | | |
| GB49375 | | |
| GB49611 | FBgn0030058 | |
| GB50282 | FBgn0036727 | |
| GB50282 | FBgn0029733 | |
| GB50282 | FBgn0039840 | |
| GB50813 | | |
| GB51260 | | |
| GB51273 | FBgn0037949 | |
| GB51367 | | |
| GB51482 | | |
| GB51556 | FBgn0259224 | |
| GB51604 | | Rx1 retinal homeobox |

| | |
|---|---|
| GB52413 | |
| GB53272 | |
| GB53972 | |
| GB54235 | |
| GB54927 | FBgn0038686 |
| GB55148 | FBgn0035688 |
| GB55764 | |
| GB55767 | |
| GB55822 | FBgn0261360 |

**Table S9.** Common putative venom constituents in Chalcidoidea parasitoid wasps *M. amalphitanum, C. solmsi, M. spermotrophus, T. pretiosum, N. vitripennis.*

| |
|---|
| NP_001155017.1 serine protease 33 precursor |
| NP_001155147.1 venom acid phosphatase-like precursor |
| NP_001155160.1 venom protein F precursor |
| NP_001155144.1 gamma-glutamyl cyclotransferase-like venom protein isoform 1 precursor |
| NP_001155153.1 aminotransferase-like venom protein 1 precursor |
| NP_001155043.1 serine protease 22 precursor |
| NP_001155086.1 glucose dehydrogenase-like venom protein |
| NP_001155148.1 carboxylesterase clade B, member 2 precursor |
| NP_001155076.1 serine protease 50 precursor |
| NP_001155015.1 serine protease precursor |
| NP_001155040.1 low-density lipoprotein receptor-like venom protein precursor |
| NP_001155145.1 gamma-glutamyl cyclotransferase-like venom protein isoform 2 |
| NP_001154998.1 cysteine-rich/KU venom protein precursor |
| NP_001155014.1 serine protease 96 precursor |
| NP_001155042.1 serine protease 97 precursor |
| NP_001155154.1 antigen 5-like protein 1 precursor |
| NP_001155016.1 serine protease homolog 29 precursor |
| NP_001155077.1 serine protease 16 precursor |
| NP_001155156.1 aminotransferase-like venom protein 2 precursor |
| NP_001154991.1 lipase A-like precursor |
| NP_001155084.1 chitinase 5 precursor |
| NP_001155079.1 serine protease homolog 42 isoform 2 precursor |
| NP_001155078.1 serine protease homolog 42 isoform 1 precursor |
| NP_001155158.1 venom laccase precursor |

| |
|---|
| NP_001155164.1 venom protein R precursor |
| NP_001155060.1 serine protease homolog 21 precursor |
| NP_001155157.1 aspartylglucosaminidase precursor |
| NP_001155159.1 laccase-like precursor |
